# A proteomics screen of HIV-1 ASP interactors reveals multifaceted effects on host cell homeostasis

**DOI:** 10.64898/2026.11.10.765637

**Authors:** Mohd. Shameel Iqbal, TaeHee Lee, Carmelo Letao Li, Yuchu Wang, Xinjie Ji, Rui Li, Xinyu Liu, Isabella Caico, Philipp Oberdoerffer, Fabio Romerio

**Author notes:** Correspondence: Department of Molecular and Comparative Pathobiology, Johns Hopkins University School of Medicine, 733 North Broadway, Baltimore, MD 21205, United States.; Department of Radiation Oncology and Molecular Radiation Sciences, Johns Hopkins University School of Medicine, 1550 Orleans Street, Baltimore, MD 21287, United States. Department of Microbiology, Heersink School of Medicine, The University of Alabama at Birmingham, Birmingham, AL, 35294, United States. LigaChem Biosciences Inc, Daejeon, Republic of Korea (TaeHee Lee). Johns Hopkins School of Medicine, Cellular and Molecular Medicine, Baltimore, MD, 21205, United States. Department of Microbiology, Department of Pediatrics, Carver College of Medicine, The University of Iowa, Iowa City, IA 52242, United States. Graduate Program in Life Sciences, University of Maryland, Baltimore, MD, 21201, United States. Brainlab, Munich, 81829, Germany (Isabella Caico).

## Abstract

Nearly four decades after the identification of the HIV-1 antisense gene *asp*, the function of its protein product remains unclear. Evidence of ASP expression *in vivo* and the high degree of conservation of the *asp* gene among pandemic HIV-1 strains suggest a beneficial role for the virus. Using a mass spectrometry (MS)-based proteomics screen for ASP interactors, we identified nuclear, cytosolic, and mitochondrial host factors, congruent with variable sub-cellular ASP localization. A majority of the host factors identified in the MS screen are involved in pathways essential for host cell homeostasis, including DNA damage repair, intracellular protein transport, mitochondrial function and protein translation. Validating the biological relevance of these findings, we demonstrate that ASP shuttles to the nucleus to promote DNA damage repair and cell viability in response to genotoxic stress. We also finds that ASP promotes translation of mitochondrially encoded host proteins involved in cellular respiration and contributes to maintain mitochondrial membrane potential. Together, these studies identify ASP as a modulator of two major pathways of cellular homeostasis in infected cells, which has direct implications for HIV-1 persistence, spread, and the long-term survival of the viral reservoir.

## Introduction

The Human Immunodeficiency Virus type 1 (HIV-1) antisense gene, *asp* was identified almost forty years ago through a search of new open reading frames in the negative strand of twelve viral isolates (*1*). Since then, the presence of *asp* has been confirmed in thousands of primary and laboratory-adapted HIV-1 strains (*2–4*). *Asp* overlaps the envelope glycoprotein gene, *env* at the Surface (SU)/Transmembrane (TM) boundary (*4, 5*). It is found in a majority of pandemic HIV-1 strains of group M, whereas it is absent in all other lentiviral species, including non-pandemic HIV-1 groups N, O, and P (*2, 4*). Within HIV-1 group M, *asp* is highly conserved among strains from high-prevalence clades and circulating recombinant forms (A, B, C, G, and CRF01_AE: ∼82% total prevalence versus D, F, H, J, K, and CRF02_AG: ∼10% total prevalence) (*4*). Primate lentiviruses outside of HIV-1 group M do not express ASP due to intermediate *stops* between the canonical start and stop codons of the predicted *asp* ORF (*2*). Conservation of the intact *asp* ORF despite constraining amino acid diversity and evolution of ENV (*3*) suggests that ASP plays a beneficial role in the HIV-1 replication cycle. Consistent with this notion, the presence of *asp* is observed at significantly higher frequency in viral isolates from people living with HIV-1 (PLWH) who progress to AIDS within 2 years of primary infection (rapid progressors) than those who progress in >20 years (long-term non-progressors) (*3*). Transcription of the *asp* gene is independent of the HIV-1 transactivator Tat, which is consistent with expression of ASP at low levels and, unlike all other viral proteins, also during latency (*6, 7*).

ASP is a protein of approximately 190 residues with a high content of hydrophobic amino acids and is predicted to contain two transmembrane domains separated by an ectodomain (*7*). The presence of cellular and humoral immune responses directed against the ASP protein in PLWH provides evidence of its expression *in vivo* during natural infection (*8–14*). We previously described a highly dynamic subcellular distribution of ASP during different phases of the virus life cycle: nuclear during viral latency and cytosolic during productive infection (*7*). Moreover, we showed that during productive infection ASP is associated with the plasma membrane as well as the envelope of infectious HIV-1 virions (*7*).

While we recently showed that the presence of ASP on the surface of infectious HIV-1 virions enhances viral fusion with the membrane of new target cells (*15*), the function of ASP within cells has not been defined. To overcome this limitation, we used mass spectrometry (MS) to identify ASP interacting partners (“ASP interactome”). This screen revealed a diverse group of host proteins that cluster around essential cellular pathways including DNA damage response and repair, intracellular protein transport, and mitochondrial function. We provide functional validation that ASP is transported to the nucleus where it responds to genotoxic stress to promote DNA repair. In addition, we show that ASP interacts with the mitochondrial protein translation machinery and contributes to stabilize mitochondrial membrane potential. These findings place ASP at the intersection of key pathways ensuring cellular homeostasis during HIV-1 infection. Our findings provide a framework for the investigation of host-virus interactions that facilitate HIV-1 persistence and the survival of infected cells.

## Results

### Construction of an HIV-1 molecular clone expressing EPEA-tagged ASP

The evolutionary mechanisms that created the *asp* gene and its high degree of conservation suggest that ASP plays an important role in the viral life cycle (*2, 3, 16*). However, the lack of known orthologs and paralogs of ASP as well as the lack of a three-dimensional structure impede an assessment of potential ASP functions. To address this limitation, we sought to isolate ASP from live cells and identify interacting factors through an unbiased mass spectrometry (MS)-based proteomics screen. The anti-ASP 324.6 antibody we describe in our previous report (*7*) is not suitable for immunoprecipitation and affinity purification applications. Thus, we modified the HIV-1JRCSF molecular clone (pYK-JRCSF) (*17*) to express a tagged ASP protein (Fig. 1A). To that end, we modified the *asp* ORF as follows: (1) we mutated codon 184 from TTA (Leu) to CCA (Pro); (2) we inserted a new codon GAA (Glu) at position 185; and (3) we mutated the next CAG codon to GCG (Ala 186). These changes created a Glutamic Acid – Proline – Glutamic Acid – Alanine (EPEA) sequence in the last four positions (aa 183-186) of ASP (Fig. 1B). This minimal epitope is targeted by the C-Tag anti-EPEA antibody in the CaptureSelect™ affinity matrix, thus allowing to capture ASP. The insertion of the EPEA tag in the *asp* ORF mutated the *env* ORF from 5’-TAC TGT AAT TCA-3’ (nt 1138-1149) to 5’-TAC <u>GCT TCT GGT</u> TCA-3’ (nt 1138-1152) resulting in a change in the amino acid sequence of ENV from ^380^YCNS^383^ to 380Y<u>ASG</u>S^384^. However, these mutations did not introduce premature *stops* in the *env* ORF and did not change any other HIV-1 sequence.

**Figure 1.**
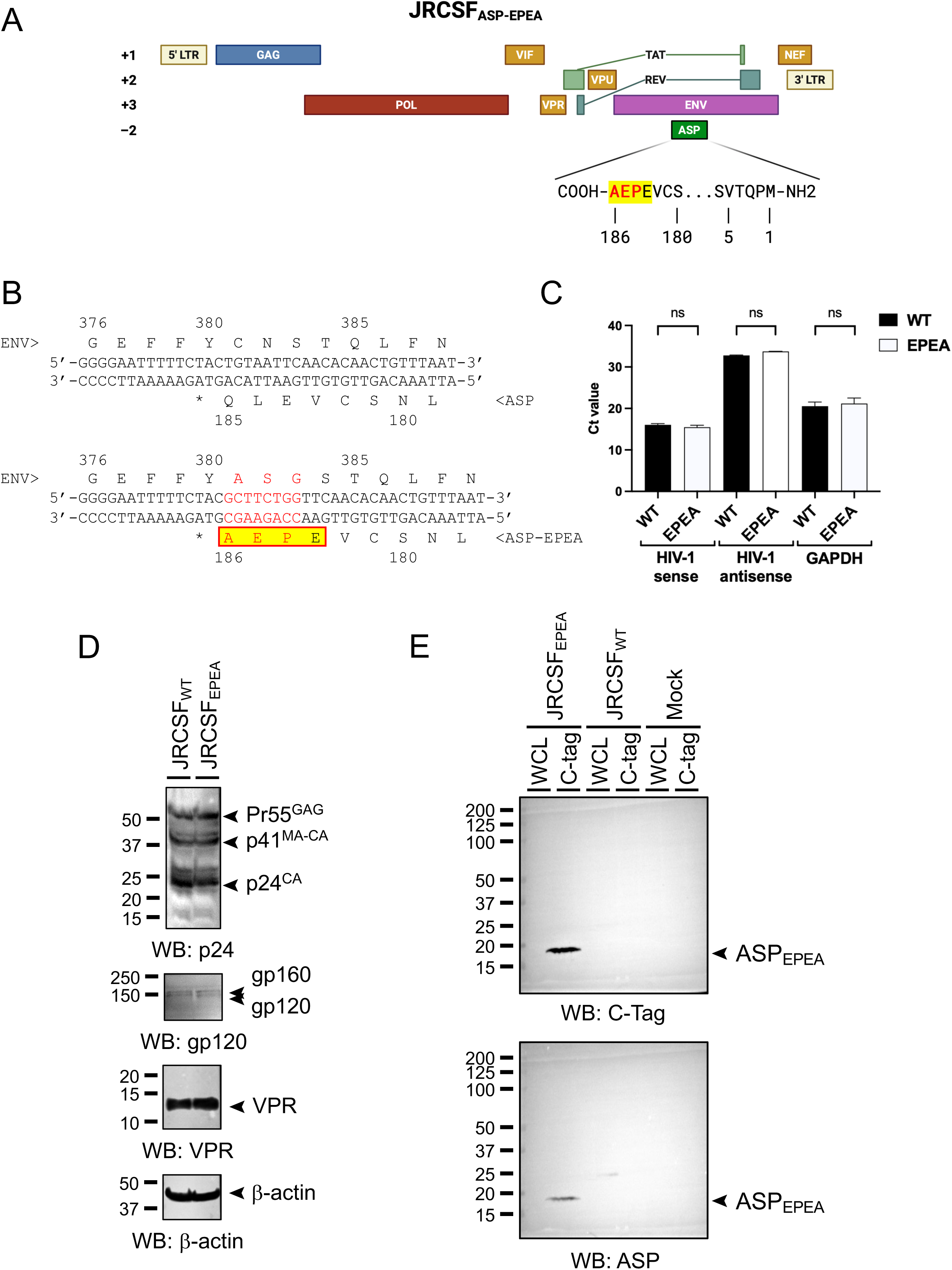
Validation of the JRCSF-based HIV-1 molecular clone expressing EPEA-tagged ASP. (A) Schema of the JRCSF molecular clone showing the position of the *asp* open reading frame. The last four C-terminal glutamic acid, proline, glutamic acid, alanine (EPEA) residues are highlighted. (B) Schema showing the genomic nucleotide sequence comprising the 3’ region of the *asp* ORF of JRCSFWT (top) and JRCSFASP-EPEA (bottom). The nucleotide substitutions that created the EPEA tag and the relative amino acid modifications are shown in red font. The EPEA tag is highlighted. (C) RT-qPCR data (Ct value) showing that the JRCSFWT (black bars) and JRCSFASP-EPEA (white bars) express the same levels of sense and antisense HIV-1 transcripts. GAPDH mRNA levels are also included as a control. (D) Western blot analyses of 293T cells transfected with the JRCSFWT and JRCSFASP-EPEA molecular clones. Whole cell lysates were analyzed with antibodies against p24^GAG^ (first blot), ENV (second blot), VPR (third blot), and β-actin (fourth blot). The two molecular clones express equal levels of HIV-1 proteins. (E) Western blot analyses of 293T cells transfected with the JRCSFWT and JRCSFASP-EPEA molecular clones. Whole cell lysates (WCE) and EPEA-enriched fractions (C-tag) were analyzed with antibodies against EPEA (C-tag; top panel) and against ASP (324.6 (*7*); bottom panel). The two antibodies detect a single band of equal molecular weight (∼20 kDa), but only in EPEA-enriched fractions from cells transfected with the JRCSFASP-EPEA molecular clone. No bands are detected in WCL or C-tag fractions from cells transfected with the JRCSFWT molecular clone or from mock-transfected cells.

First, we verified that the HIV-1 molecular clone expressing the EPEA-tagged ASP expressed the same levels of HIV-1 sense and antisense transcripts. To that end, we transfected the wildtype (JRCSFWT) and the EPEA-modified (JRCSFASP-EPEA) molecular clones into HEK293T cells. We then used total RNA to perform RT-qPCR analyses using primers specific for sense and antisense HIV-1 transcripts (*18, 19*). As a control, we measured the expression levels of GAPDH transcripts. The two molecular clones expressed identical levels of sense and antisense transcripts (Fig. 1C). Next, we prepared whole cell lysates (WCL) from HEK293T cells transfected with the JRCSFWT or JRCSFASP-EPEA molecular clones and we performed Western blot (WB) analyses to evaluate expression levels of GAG, VPR, and ENV proteins. As a control, we performed WB with antibodies directed against β-actin. Again, we confirmed that the two molecular clones express similar levels of HIV-1 proteins (Fig. 1D). Moreover, these finding establish that the mutations in *env* do not alter ENV expression or stability.

Finally, we assessed whether the CaptureSelect™ affinity matrix captures ASP-EPEA efficiently and selectively. Thus, we performed WB analyses using either WCL or CaptureSelect™-enriched fractions (EPEA) from HEK293T cells transfected with the JRCSFWT or JRCSFASP-EPEA molecular clones. As a control, we used mock-transfected cells. A band of the expected molecular weight (∼22 kDa) was readily detectable with both anti-EPEA C-Tag (Fig. 1E) and anti-ASP 324.6 mAbs (Fig. 1F) in the CaptureSelect™-enriched fraction from cells transfected with the JRCSFASP-EPEA molecular clone. We did not observe ASP signal in WCL from HEK293T cells transfected with either molecular clone and probed with either antibody, or in the CaptureSelect™-enriched fraction from cells transfected with the JRCSFWT molecular clone (Fig. 1E and F). Detection of ASP only in CaptureSelect™-enriched fractions was expected because the negative sense promoter in the HIV-1 3’ LTR that drives ASP expression has weak transcriptional activity (*6, 19*). Finally, no band was visible in WB with mock-transfected cells under any of the conditions described above (Fig. 1E and F).

Altogether, the insertion of the EPEA tag did not impact sense and antisense HIV-1 transcription nor the expression of viral proteins (Fig. 1C and D). The lack of bands other than ASP in the CaptureSelect™-enriched fraction from cells transfected with the JRCSFASP-EPEA molecular clone after probing with C-Tag antibodies indicates that the EPEA tag allows to capture ASP in a highly specific manner (Fig. 1E). Finally, the ability to detect ASP-EPEA in the CaptureSelect™-enriched fraction but not WCL from cells transfected with the JRCSFASP-EPEA molecular clone after probing with either anti-EPEA C-Tag (Fig. 1E) or anti-ASP 324.6 mAbs (Fig. 1F) indicates that the EPEA tag allows to efficiently capture ASP.

### Mass spectrometry-based screen of ASP interacting partners

After establishing and validating the JRCSFASP-EPEA molecular clone, we proceeded with the proteomics screen of ASP interacting partners (Fig. 2A). We transfected the wildtype and EPEA-modified HIV-1 molecular clones into HEK293T cells. After 72 hours we prepared WCL, which were passed through a C-Tag CaptureSelect™ affinity matrix. After washing, bound proteins were eluted, and an aliquot of the two eluates were analyzed by SDS-PAGE. The results in Fig. 2B show qualitative and quantitative differences between the eluates from cells transfected with the JRCSFWT or JRCSFASP-EPEA molecular clones. After digesting the recovered proteins with trypsin, 500 and 1500 ng of desalted peptides from each sample were analyzed by liquid chromatography/tandem mass spectrometry (LC-MS/MS). This procedure was performed with samples from two independent transfection experiments. Analysis of the MS results found 431 proteins significantly enriched or depleted in the ASP-EPEA sample in both biological replicates (linFC ≥ 2.0 or linFC ≤ -2.0, respectively; Data S1) compared to the WT samples (*p <* 0.05) (Fig. 2C): 376 proteins were enriched in the ASP-EPEA samples (Fig. S1) and 55 in the WT samples (Fig. S2). Normalization was performed based on proteins represented in equal amounts in the ASP and ASP-EPEA samples.

**Figure 2.**
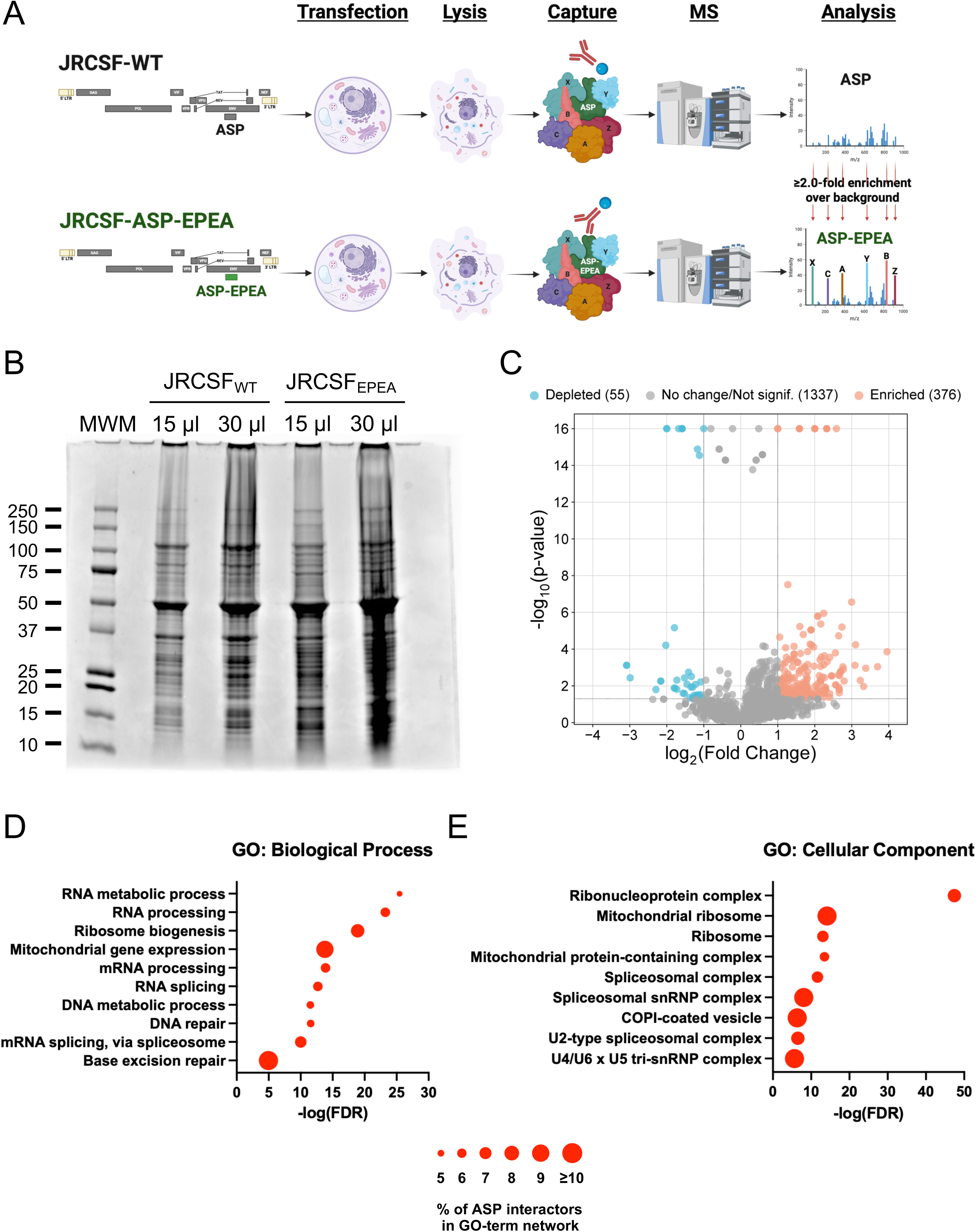
Mass spectrometry-based identification of ASP interacting partners. (A) Schema of the experimental approach. The JRCSFWT and JRCSFASP-EPEA molecular clones are first transfected into 293T cells. Whole cell lysates are then incubated with CaptureSelect™ affinity matrix. The recovered material is digested with trypsin and the peptides analyzed by mass spectrometry to identify proteins significantly enriched in 293 T cells transfected with JRCSFASP-EPEA compared to cells transfected with JRCSFWT. The experiment was performed with samples from two independent biological replicates. (B) CaptureSelect™-enriched fractions (15 and 30 μl) from 293T cells transfected with JRCSFWT or JRCSFASP-EPEA molecular clones were separated on a 4-20% Tris-Glycine polyacrylamide gel. Both qualitative and quantitative differences are visible between samples from cells transfected with the two molecular clones. (C) Volcano plot analysis of proteins identified by mass spectrometry in both biological replicates. Proteins significantly enriched in samples from cells transfected with the JRCSFASP-EPEA molecular clone compared to cells transfected with the JRCSFWT molecular clone (376 proteins) are shown in shades of pink. Vice versa, proteins significantly enriched in samples from cells transfected with JRCSFWT compared to cells transfected with JRCSFASP-EPEA (55 proteins) are shown in shades of aqua. Proteins not-significantly enriched in either sample (1337 proteins) are shown in gray. (D) Gene Ontology (GO) Biological Process terms significantly enriched among proteins enriched in JRCSFASP-EPEA samples compared to JRCSFWT samples. (E) GO Cellular Component terms significantly enriched in JRCSFASP-EPEA samples compared to JRCSFWT samples. Enrichment analysis was performed using the STRING Protein-Protein Interaction Networks Functional Enrichment Analysis tool, and terms meeting a -log10(FDR) ≥ 5 threshold are shown. Dot size represents the percentage of proteins in the term found to be ASP interactors.

Analysis of the MS results using the STRING database of protein-protein interactions revealed that more than half of the proteins enriched in the ASP-EPEA samples (178 of 341) belong to only three primary and two secondary functional pathways (Table 1; Fig. 2D and E). First, the MS screen revealed a major cluster of cellular proteins with a role in DNA replication stress, DNA repair, and chromatin remodeling (Fig. S3). Notably, the ASP interactome included proteins involved in all major DNA repair pathways: mismatch repair (MMR), base excision repair (BER), nucleotide excision repair (NER), non-homologous end-joining (NHEJ), homologous recombination (HR), and the replication stress response (*20–22*) (Table 2). Second, we found a very high enrichment of proteins involved in RNA metabolism and spliceosome-associated RNA processing, including core spliceosome factors, RNA helicases, exon definition factors, RNA maturation and polyadenylation factors, RNA structure and surveillance factors (Fig. S3). STRING analysis showed that both the DNA and RNA metabolism sets are intimately and functionally connected with each other, as evidenced by multiple edges linking proteins in one subset with proteins in the other subsets (Fig. S3). Third, we identified a number of proteins involved in ribosome biogenesis, translation-associated stress regulation, mitochondrial ribosome proteins, and translation initiation and elongation factors (Table 1; Fig. S4). In addition to these three major functional pathways, we uncovered proteins associated with two secondary pathways. The first one is a nucleocytoplasmic trafficking signature that includes karyopherins, importins, and exportins (Table 1; Fig. S4), and the second is a cytoskeletal and endomembrane trafficking signature that includes actin-associated proteins, myosins, spectrins, coatomer I (COPI) subunits, and clathrin/endocytosis components (Table 1; Fig. S4). In all of these cases, the number of edges (known or predicted protein-protein interactions) that link the proteins within the network (nodes) was significantly larger than what would be expected for a sample of equal size and degree of distribution drawn randomly from the human proteome (*p* < 10^-16^). On the other hand, the 55 proteins that were depleted in the ASP-EPEA vs. WT dataset comprise a large number of keratin/epithelial contaminants and abundant background proteins (Fig. S2). Therefore, the MS screen identified multiple potential ASP interacting partners. In many cases, these proteins are biologically connected and are known to play essential cellular functions.

**Table 1.**
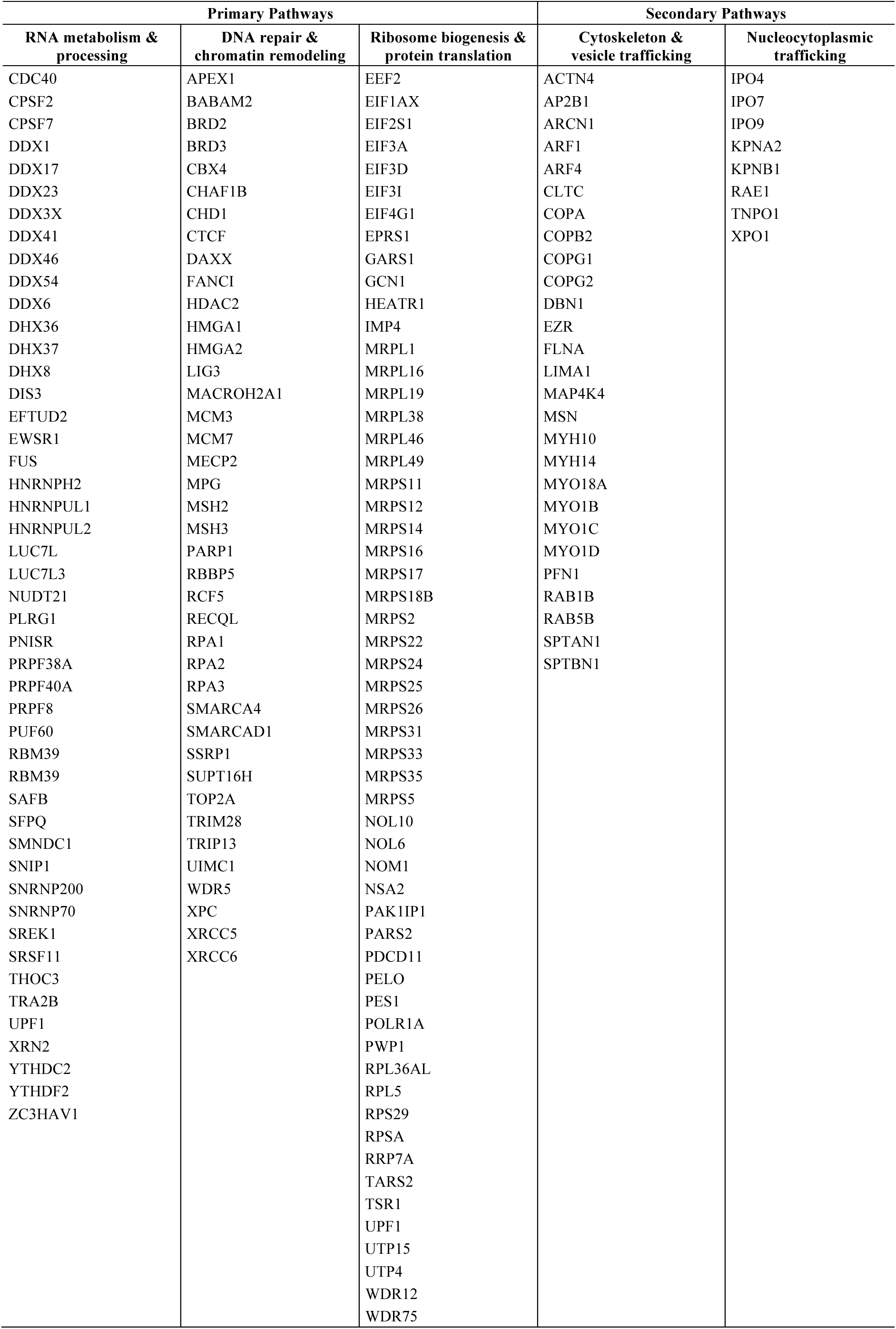
Selected list of host factors (178 of 341) identified in the ASP interactome through IP-MS. See Data S1 for complete list.

| **Primary Pathways** |  |  | **Secondary Pathways** |  |
|---|---|---|---|---|
| **RNA metabolism & processing** | **DNA repair & chromatin remodeling** | **Ribosome biogenesis & protein translation** | **Cytoskeleton & vesicle trafficking** | **Nucleocytoplasmic trafficking** |
| CDC40 | APEX1 | EEF2 | ACTN4 | IPO4 |
| CPSF2 | BABAM2 | EIF1AX | AP2B1 | IPO7 |
| CPSF7 | BRD2 | EIF2S1 | ARCN1 | IPO9 |
| DDX1 | BRD3 | EIF3A | ARF1 | KPNA2 |
| DDX17 | CBX4 | EIF3D | ARF4 | KPNB1 |
| DDX23 | CHAF1B | EIF3I | CLTC | RAE1 |
| DDX3X | CHD1 | EIF4G1 | COPA | TNPO1 |
| DDX41 | CTCF | EPRS1 | COPB2 | XPO1 |
| DDX46 | DAXX | GARS1 | COPG1 |  |
| DDX54 | FANCI | GCN1 | COPG2 |  |
| DDX6 | HDAC2 | HEATR1 | DBN1 |  |
| DHX36 | HMGA1 | IMP4 | EZR |  |
| DHX37 | HMGA2 | MRPL1 | FLNA |  |
| DHX8 | LIG3 | MRPL16 | LIMA1 |  |
| DIS3 | MACROH2A1 | MRPL19 | MAP4K4 |  |
| EFTUD2 | MCM3 | MRPL38 | MSN |  |
| EWSR1 | MCM7 | MRPL46 | MYH10 |  |
| FUS | MECP2 | MRPL49 | MYH14 |  |
| HNRNPH2 | MPG | MRPS11 | MYO18A |  |
| HNRNPUL1 | MSH2 | MRPS12 | MYO1B |  |
| HNRNPUL2 | MSH3 | MRPS14 | MYO1C |  |
| LUC7L | PARP1 | MRPS16 | MYO1D |  |
| LUC7L3 | RBBP5 | MRPS17 | PFN1 |  |
| NUDT21 | RCF5 | MRPS18B | RAB1B |  |
| PLRG1 | RECQL | MRPS2 | RAB5B |  |
| PNISR | RPA1 | MRPS22 | SPTAN1 |  |
| PRPF38A | RPA2 | MRPS24 | SPTBN1 |  |
| PRPF40A | RPA3 | MRPS25 |  |  |
| PRPF8 | SMARCA4 | MRPS26 |  |  |
| PUF60 | SMARCAD1 | MRPS31 |  |  |
| RBM39 | SSRP1 | MRPS33 |  |  |
| RBM39 | SUPT16H | MRPS35 |  |  |
| SAFB | TOP2A | MRPS5 |  |  |
| SFPQ | TRIM28 | NOL10 |  |  |
| SMNDC1 | TRIP13 | NOL6 |  |  |
| SNIP1 | UIMC1 | NOM1 |  |  |
| SNRNP200 | WDR5 | NSA2 |  |  |
| SNRNP70 | XPC | PAK1IP1 |  |  |
| SREK1 | XRCC5 | PARS2 |  |  |
| SRSF11 | XRCC6 | PDCD11 |  |  |
| THOC3 |  | PELO |  |  |
| TRA2B |  | PES1 |  |  |
| UPF1 |  | POLR1A |  |  |
| XRN2 |  | PWP1 |  |  |
| YTHDC2 |  | RPL36AL |  |  |
| YTHDF2 |  | RPL5 |  |  |
| ZC3HAV1 |  | RPS29 |  |  |
|  |  | RPSA |  |  |
|  |  | RRP7A |  |  |
|  |  | TARS2 |  |  |
|  |  | TSR1 |  |  |
|  |  | UPF1 |  |  |
|  |  | UTP15 |  |  |
|  |  | UTP4 |  |  |
|  |  | WDR12 |  |  |
|  |  | WDR75 |  |  |

**Table 2.** DNA repair factors interacting with ASP.

| **Protein** | **Lin(FC)** | **p-value** | **Pathway** | **Function** |
|---|---|---|---|---|
| MSH3 | 12.96 | 0.0009 | MMR | MutSβ subunit; Mismatch sensor of large indels |
| RPA1 | 10.38 | 0.0009 | Multiple | 70 kDa subunit of RPA; ssDNA binding, scaffolding |
| MPG | 7.97 | <0.0001 | BER | Short-patch repair; Base damage sensor |
| MSH2 | 6.83 | 0.0012 | MMR | MutSβ subunit; Mismatch sensor of large indels |
| LIG3 | 6.77 | 0.0009 | BER; SSB | Short-patch repair; DNA Ligase |
| RECQL | 5.48 | 0.0028 | BER; DSB | Long-patch repair; DNA helicase |
| RPA2 | 5.29 | 0.0028 | Multiple | 32 kDa subunit of RPA; ssDNA binding, scaffolding |
| XPC | 5.00 | 0.0001 | NER | Global genome repair; DNA damage sensor |
| RPA3 | 4.53 | 0.0003 | Multiple | 14 kDa subunit of RPA; ssDNA binding, scaffold |
| APEX1 | 3.87 | 0.0377 | BER | Short and long-patch repair; Endonuclease |
| PARP1 | 3.77 | <0.0001 | BER; SSB | SSB sensor; scaffold |
| FANCI | 3.16 | 0.0011 | DSB | Subunit of Fanconi anemia complex I; Replication fork stabilizer |
| XRCC5 | 3.02 | <0.0001 | DSB | Subunit of KU heterodimer; DNA break stabilizer |
| BRCC45 | 3.00 | <0.0001 | DSB | Subunit of BRCA1 complex; Promotes NHEJ |
| UIMC1 | 3.00 | <0.0001 | DSB | Subunit of BRCA1 complex; Promotes NHEJ |
| XRCC6 | 2.49 | 0.0004 | DSB | Subunit of KU heterodimer; DNA break stabilizer |

BER: base excision repair; MMR: mismatch repair; NER: nucleotide excision repair; DSB: double strand break; SSB: single strand break.

### ASP protects from genotoxic stress

Among the ASP interactors, we identified key components of several repair pathways targeting both single-and double-stranded DNA lesions (Table 2). A subset of ASP:repair factor interactions were validated in 293T cells using co-IP with antibodies specific for DNA repair factors followed by WB for FLAG-tagged ASP protein (Fig 3A). Reverse IP confirmed ASP interaction with RPA32 and RPA70, two components of the RPA ssDNA binding protein complex involved in multiple DNA repair pathways, including HR, the replication stress response and long-patch BER (Fig 3B, C). No interaction was observed with GFP fused to a nuclear localization signal and three copies of the FLAG epitope (GFP-NLS-3×FLAG) (Fig. 3B, C). Based on ASP association with four central BER factors, PARP1, LIG3 APEX1 and RPA (Table 1), we focused our functional validation efforts on the repair of oxidative ssDNA base lesions, a common source of genotoxic stress in both dividing and resting cells. To assess whether ASP alters the accumulation of oxidative damage, we measured the turnover of the most prevalent H2O2-induced DNA lesion, 8-oxo-Guanine (8-oxo-G), in the presence or absence of ASP protein. HeLa cells stably expressing ASP showed comparable initial 8-oxo-G accumulation, indicating efficient DNA damage induction. Following a 4-hour recovery phase, ASP-expressing cells displayed a sharp drop in 8-oxo-G compared to control cells transduced with empty vector (Fig. 3D). Consistent with this, we observed improved resolution of ssDNA breaks, as determined using the alkaline Comet assay, demonstrating a role for ASP in promoting BER (Fig. 3E). Improved repair was associated with a decrease in sensitivity to H2O2-induced cell death (Fig. 3F). A similar effect was observed for a second DNA base-damaging agent, methyl-methane sulfonate (MMS), corroborating the DNA damage-protective role for ASP in response to DNA lesions repaired by BER (Fig. S5).

**Figure 3.**
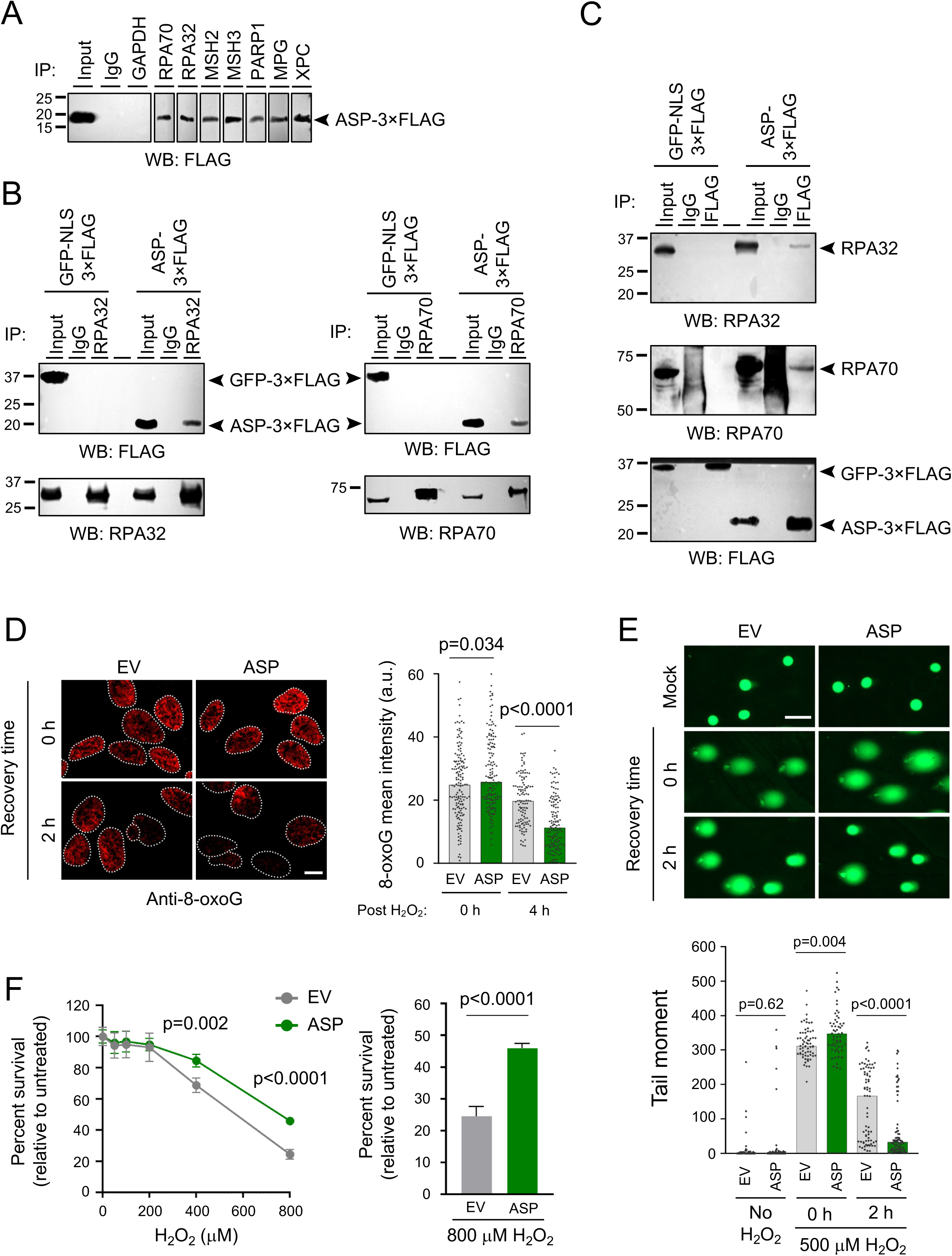
ASP protects from oxidative DNA damage. (A) Anti-FLAG Western blot of cellular lysates before (input) and after co-immunoprecipitation (co-IP) with antibodies for the indicated proteins in 293T cells transfected with ASP-3×FLAG expression vector. (B) Western blot with anti-FLAG antibodies (top blots) before (input) and after anti-RPA32 (top left), anti-RPA70 (top right), or IgG co-IP in 293T cells transiently expressing FLAG-tagged GFP or ASP protein. Bottom blots show efficient recovery of RPA32 and RPA70 from both cell lines after IP with the respective antibodies. (C) Western blot with anti-RPA32 (top blot) or anti-RPA70 antibodies (middle blot) before (input) and after co-IP with anti-FLAG or IgG antibodies in 293T cells transiently expressing FLAG-tagged GFP or ASP protein. Bottom blot shows efficient recovery of ASP and GFP from the respective cell lines after IP with anti-FLAG antibodies. (D) Immunofluorescence analysis of 8-oxo-G lesion abundance in HeLa cells stably transduced with empty vector (EV) or ASP-3×FLAG at the indicated time points after H2O2 treatment (500 µM, 30 min). Bee-swarm plots depict mean nuclear intensities, and bars indicate the median (n = 158, 143, 115 and 133 cells, respectively). One of two independent experiments is shown. (E) Alkaline Comet assay in HeLa cells as in (D), at the indicated time points after H2O2 treatment or left untreated (no H2O2). Bee-swarm plots depict Olive tail moment, and bars indicate the median (n = 87, 83, 65, 57, 73 and 81 cells, respectively). P values are based on two-sided Mann-Whitney U test. (F) Cell viability assay (MTT) in HeLa cells as in (D) exposed to the indicated doses of H2O2 for 24 h, followed by 24 h recovery. Values are normalized to no treatment and expressed as mean and SD (n = 4 technical replicates), bar chart depicts values at 800 µM H2O2. P values are based on Student’s two-tailed t-test. One of two independent experiments is shown.

Notably, ASP-mediated enhancement of DNA repair extends to at least one additional repair pathway, the repair of UV lesions, which requires the NER mediator and ASP interactor, XPC. ASP suppressed both UV-induced DNA break accumulation and cell death, supporting the notion that ASP is a more general effector of DNA repair responses (Fig. S6). How interaction with ASP enhances repair factor function will be an important avenue of future investigation.

### ASP interacts with chromatin factors involved in DNA repair

Chromatin context directly affects DNA repair factor recruitment (*23*), and all ASP-interacting host factors involved in chromatin organization identified by MS are involved in DNA repair (Fig. 4A, Table 3) (*23, 24*). We thus asked if ASP, like the DNA repair machinery, is directly associated with chromatin. First, we confirmed the interaction between ASP and two chromatin remodeling factors, SSRP1 and SUPT16H, by two-way co-IP (Fig. 4B, C). We further observed an enrichment of core histones in the chromatin fraction of cells expressing ASP compared to cells expressing an NLS-containing GFP protein (Fig. 4D), which points to a role for ASP in promoting a more condensed chromatin state. Supporting this notion, ASP-expressing cells showed increased resistance to MNase I digestion (Fig. 4E-H and S7). Of note, we and others have previously identified dynamic chromatin compaction as a means to control DNA repair outcomes (*25–27*). Among the histones enriched in the chromatin fraction upon ASP expression were macroH2A1 isoforms, which are involved in heterochromatin formation, genome maintenance and both positive and negative regulation of gene expression (*24, 27–32*). Of relevance for ASP function in BER, work from our lab and others has shown a role for macroH2A1 in repair processes associated with the removal of oxidative lesions (*33, 34*). We did not observe a change in chromatin association in response to H2O2 or UV treatment (Fig. S8), suggesting that ASP establishes a chromatin environment that aids repair, rather than being recruited to chromatin in response to damage. This is consistent with the breadth of ASP-interacting chromatin components. However, we cannot rule out that other sources of damage may alter the interaction of ASP with chromatin.

**Figure 4.**
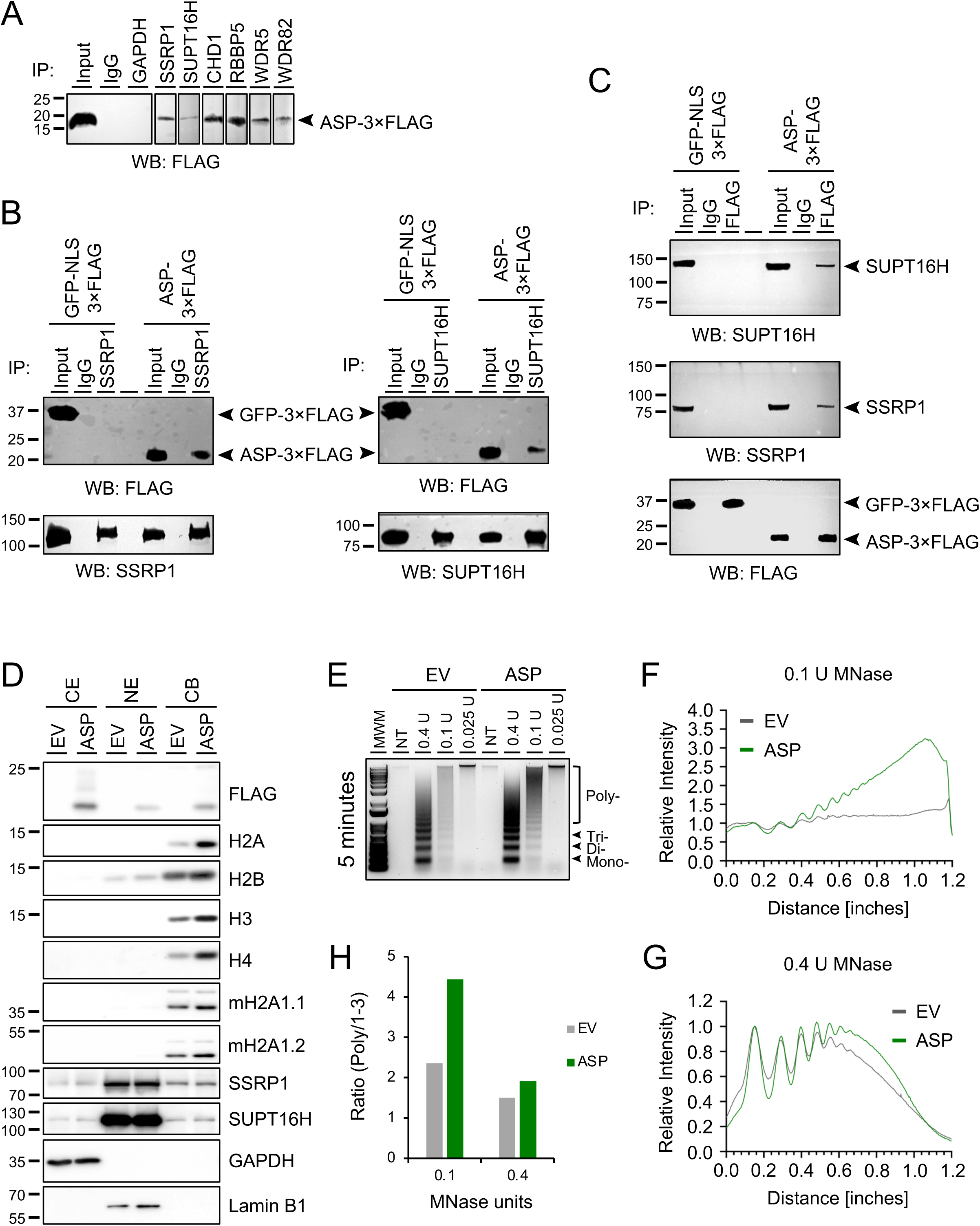
ASP is a chromatin interactor and promotes chromatin compaction. (A) Anti-FLAG Western blot of cellular lysates before (input) and after co-immunoprecipitation (co-IP) with antibodies for the indicated proteins in 293T cells transfected with ASP-3×FLAG expression vector. (B) Western blot with anti-FLAG antibodies (top blots) before (input) and after anti-SSRP1 (top left), anti-SUPT16H (top right), or IgG co-IP in 293T cells transiently expressing FLAG-tagged GFP or ASP protein. Bottom blots show efficient recovery of SSRP1 and SUPT16H from both cell lines after IP with the respective antibodies. (C) Western blot with anti-SSRP1 (top blot) or anti-SUPT16H antibodies (middle blot) before (input) and after co-IP with anti-FLAG or IgG antibodies in 293T cells transiently expressing FLAG-tagged GFP or ASP protein. Bottom blot shows efficient recovery of ASP and GFP from the respective cell lines after IP with anti-FLAG antibodies. (D) Expression of ASP-3×FLAG and the indicated proteins in cytoplasmic extract (CE), soluble nuclear extract (NE) and chromatin bound protein (CB) from HeLa cells stably transduced with empty vector (EV) or ASP-3×FLAG. One of three independent experiments is shown. (E) nucleosome profiles from HeLa cells as in (D) treated for 5 min with MNase at the indicated concentrations. MWM: molecular weight marker, U: unit. Mono, Di and Tri-nucleosome are highlighted. (F, G) Nucleosome intensity profiles of samples in (E) for the indicated MNase concentrations, normalized to the mono-nucleosome signal. Distance measures start at the bottom the gel (mono-nucleosome). (H) Ratio of poly-nucleosomes over mono- to tri-nucleosomes for cell lines form (D) at the indicated MNase concentrations. Consistent results were obtained in an independent experiment following 20 minutes of MNase treatment (**Fig. S7**).

**Table 3.** Chromatin factors involved in DNA repair interacting with ASP.

| Protein | Lin(FC) | p-value | Complex | Role in DNA damage repair |
|---|---|---|---|---|
| SUPT16H | 9.99 | 0.0108 | FACT | TC-BER, NER, SSB and DSB repair |
| RBBP5 | 4.99 | 0.0159 | SET1A/B | Protects genome integrity during replication stress |
| CHD1 | 4.73 | <0.0001 | – | DSB repair through HR |
| SSRP1 | 3.94 | 0.0009 | FACT | TC-BER, NER, SSB and DSB repair |
| CHAF1B | 3.87 | 0.0338 | CAF-1 | DSB repair and NER |
| WDR82 | 3.87 | 0.0338 | SET1A/B | Protects genome integrity during replication stress |
| WDR5 | 3.72 | <0.0001 | SET1A/B | Protects genome integrity during replication stress |
| NAP1L1 | 3.54 | 0.0243 | – | DSB repair through HR |
| CTCF | 3.31 | 0.0009 | – | DSB repair |
| MacroH2A1 | 3.22 | 0.0252 | – | DSB repair; replication stress |
| SMARCA4 | 2.45 | 0.0443 | BAF | DSB repair (HR and NHEJ) |
| SMARCAD1 | 2.45 | 0.0476 | – | SSB and DSB repair |

### Mechanisms of ASP nuclear import

Nuclear localization of proteins produced in the cytoplasm relies on nuclear import pathways. The MS screen of ASP interactors identified 3 karyopherins (Kapα2/KPNA2, Kapβ1/KPNB1, Kapβ2/TNPO1) and 3 importins (IPO4/RanBP4, IPO7/RanBP4, IPO9/RanBP4) (Table 1). We validated a subset of the interactions using co-IP against FLAG-tagged ASP protein (Fig 5A). Next, we studied the relevance of such interactions in the mechanisms of ASP nuclear import, an important independent confirmation of the detection of ASP within the nucleus (*7*) and a prerequisite for its function in DNA repair and chromatin organization. The karyopherins, Kapα2 and Kapβ1 function cooperatively: the former binds cargo carrying a canonical nuclear localization signal (cNLS consensus: K-K/R-X-K/R, where X is any residue), and the latter transports the complex into the nucleus. The karyopherin, TNPO1, on the other hand, performs both functions: it binds the cargo and transports it into the nucleus. TNPO1 recognizes a non-canonical proline-tyrosine NLS (PY-NLS) rich in hydrophobic residues (hPY-NLS consensus: Φ-G/A/S-Φ-Φ-X6-13-R/K/H-X2-5-P-Φ, where Φ is a hydrophobic residue, and X is any residue). ASP contains a KNPRNK motif (aa 51-56) that resembles but does not closely match the cNLS consensus sequence. However, ASP contains three motifs (aa 41-59, aa 96-111, aa 116-134; Fig. 5B) that closely match the hPY-NLS consensus sequence. To assess whether these motifs are involved in the interaction of ASP with TNPO1, we replaced key residues in the ASP hPY-NLS motifs with Alanine (ASP-ΔPY-NLS; Fig. 5B) and then we performed co-IP with TNPO1 antibodies (Fig. 5C). ASP-ΔPY-NLS failed to interact with TNPO1 (Fig. 5C), which led to reduced nuclear localization of ASP-ΔPY-NLS compared to wildtype ASP (Fig. 5D). This confirms that nuclear import of ASP relies on TNPO1, which emerged as an ASP-interacting factor in our MS analysis. It will be interesting to investigate how ASP nuclear import is regulated in response to DNA damage and other sources of cellular stress.

**Figure 5.**
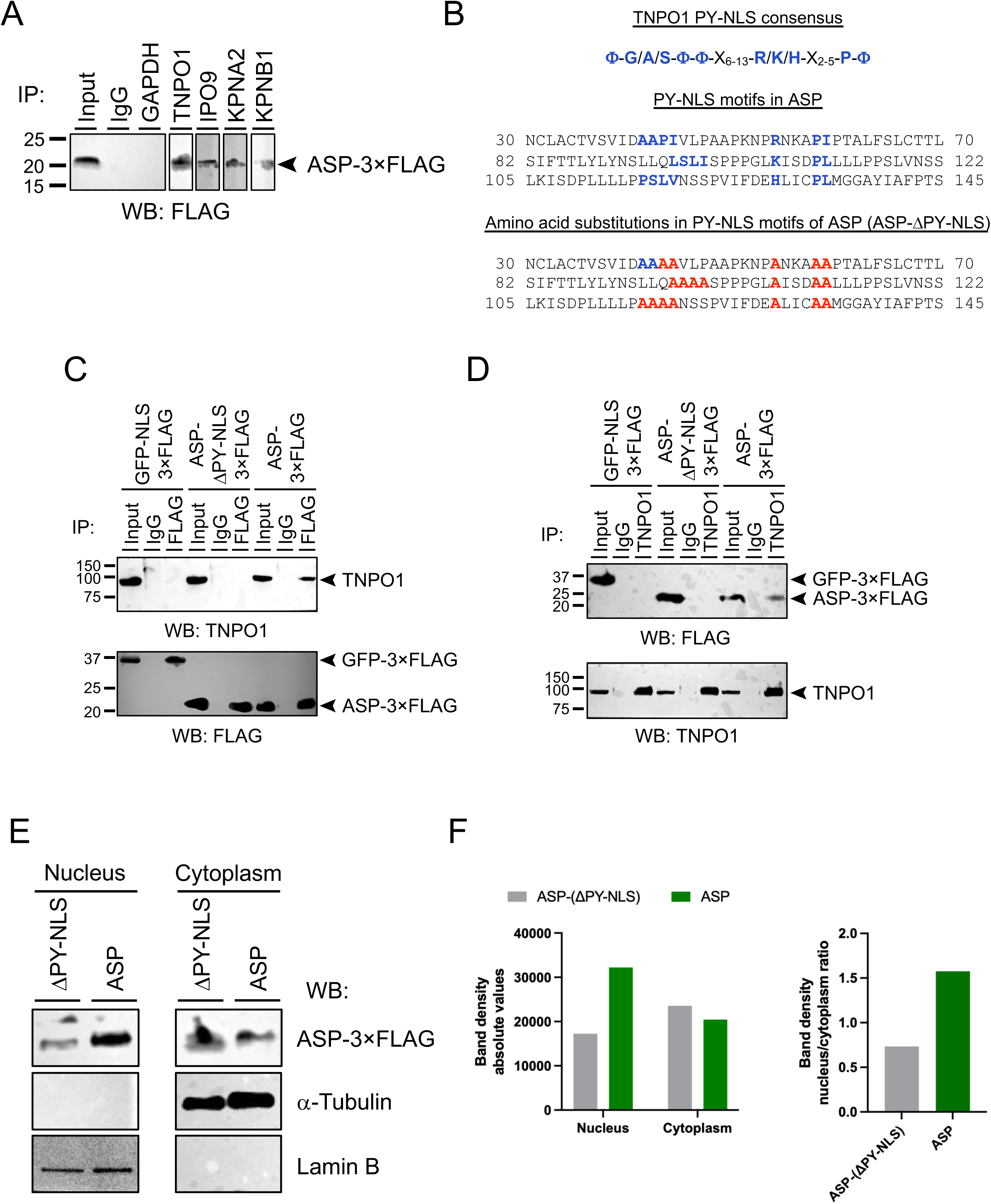
ASP shuttles into the nucleus via interaction with TNPO1 karyopherin. (A) Validation of the interaction between ASP and four karypherins/importins (TNPO1, KPNA2, KPNB1, IPO9). 293T cells were transfected with a vector expressing ASP-3×FLAG. Whole cell lysates were incubated with antibodies specific for the proteins above followed by immunoprecipitation with Protein A/G magnetic beads. Immunocomplexes were analyzed in Western blot with anti-FLAG antibodies. (B) Amino acid sequence of three regions of ASP containing motifs with similarity to the TNPO1 consensus recognition sequence (Φ-G/A/S-Φ-Φ-X6-13-R/K/H-X2-5-P-Φ). Residues in blue (top) indicate positions within the motif that are critical for interaction with TNPO1. Residues in red (bottom) indicate amino acid substitutions introduced in the ASP sequence to generate the ASP-ΔPY-NLS mutant and that are expected to disrupt the interaction with TNPO1. (C) and (D) Two-way co-immunoprecipitation analyses of the interaction between TNPO1 and ASP-3×FLAG, ASP-ΔPY-NLS-3×FLAG, or GFP-NLS-3×FLAG. In (C), whole cell lysates from 293T cells transfected with each of the three expression vectors were immunoprecipitated with anti-FLAG antibodies, and the immunocomplexes were then analyzed in Western blot with antibodies against TNPO1 (top blot) or with FLAG (bottom blot). In the reverse experiment (D), cell lysates were immunoprecipitated with antibodies anti-TNPO1 antibodies, and the immunocomplexes were then analyzed in Western blot with antibodies against FLAG (top blot) or with TNPO1 (bottom blot). Amino acid substitutions in the ASP motifs that closely match the TNPO1 consensus recognition sequence abolish the interaction between ASP and TNPO1. (E) Western blot analysis of cytoplasm and nuclear fractions from 293T cells transfected with ASP-3×FLAG or ASP-ΔPY-NLS-3×FLAG vectors using anti-FLAG antibodies. Disruption of the TNPO1 recognition motifs in the ASP sequence diminish nuclear levels of ASP. Analysis with anti-tubulin and anti-lamin B antibodies confirms efficient recovery of nuclear and cytoplasmic fractions. (F) The graph on the left shows quantification of ASP band intensities in the nuclear and cytoplasm fractions analyzed with anti-FLAG antibodies in the Western blot in (E). Band intensities were quantified using an iBright 1500 gel documentation system (Thermo Fisher). The graph on the right report the ratio of nuclear/cytoplasm band intensities for cells transfected with each of the two vectors.

### Effect of ASP on mitochondrial membrane potential

Among the ASP interactors involved in protein translation, we identified 21 proteins that constitute the large and small subunits of the mitochondrial ribosomes (mitoribosomes) (Table 1), which contain approximately 80 proteins. After validating the binding of ASP with some of the 21 proteins identified in the MS screen (Fig. 6A), we assessed whether the interaction of ASP with mitoribosomes might affect the translation and expression levels of proteins encoded in the mitochondrial genome (mtDNA). In vertebrates, the mitochondrial genome encodes 13 proteins, all of which are subunits of mitochondrial complexes I, III, IV, and V involved in the electron transport chain that generates cellular energy in the form of ATP. Thus, we compared the expression levels of 6 of the 13 mtDNA-encoded proteins: ND3 (Complex I), Cyt-B (Complex III), COX1 and COX2 (Complex IV), ATP6 and ATP8 (Complex V) in cells transfected with an ASP-expressing vector versus an empty vector. Cells expressing ASP showed higher (∼1.5- to ∼2.5-fold) levels of all 6 proteins tested (Fig. 6B, C).

**Figure 6.**
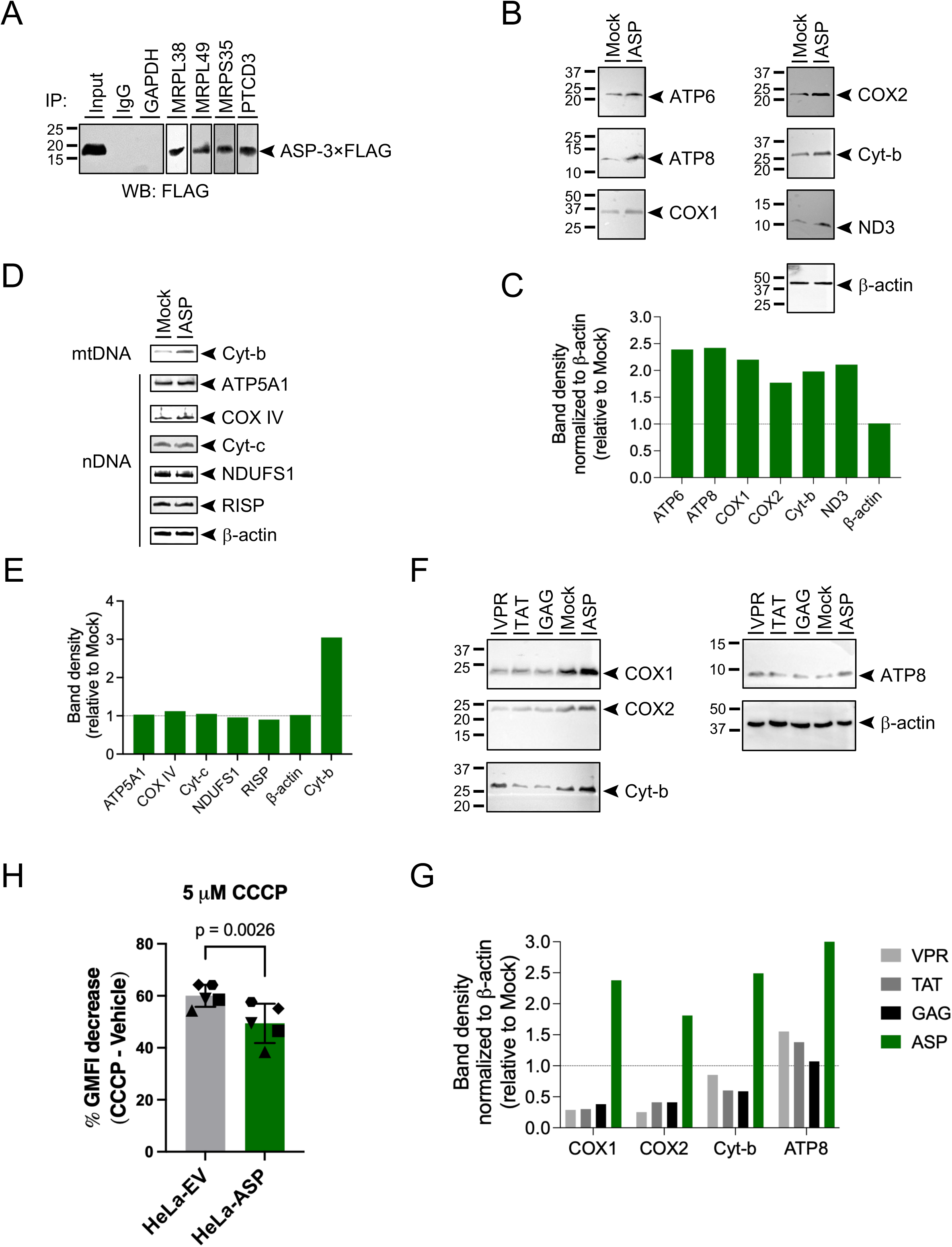
ASP maintains mitochondrial membrane potential in response to depolarizing agents. (A) Validation of the interaction between ASP and four mitochondrial ribosome proteins: MRPL38 and MRPL49 (large ribosomal subunit), MRPS35 and PTCD3/MRPS39 (small ribosomal subunit). 293T cells were transfected with a vector expressing ASP-3×FLAG. Whole cell lysates were incubated with antibodies specific for the proteins above followed by immunoprecipitation with Protein A/G magnetic beads. Immunocomplexes were analyzed in Western blot with anti-FLAG antibodies. (B) Expression levels of six mtDNA-encoded proteins (ATP6, ATP8, COX1, COX2, Cytochrome B, ND1) in 293T cells transfected with a vector expressing ASP-3×FLAG or with an empty vector (mock). As a control, lysates were also analyzed with antibodies against β-actin. (C) Intensity of the bands in panel B were determined with a Thermo Fisher iBright 1500 Imaging System and plotted as the ratio of ASP-expressing over control cells. (D) Expression levels of five nDNA-encoded mitochondrial respiratory complexes (ATP5A1, COX IV, Cytochrome C, NDUFS1, RISP) in 293T cells transfected with a vector expressing ASP-3×FLAG or with an empty vector (mock). As a control, lysates were also analyzed with antibodies against β-actin and the mtDNA-encoded protein, Cytochrome B. (E) Intensity of the bands in panel D were determined with a Thermo Fisher iBright 1500 Imaging System and plotted as the ratio of ASP-expressing over control cells. (F) Expression levels of four mtDNA-encoded proteins (ATP8, COX1, COX2, Cytochrome B) in 293T cells transfected with vectors expressing HIV-1 VPR, GAG, TAT, ASP-3×FLAG, or with an empty vector (mock). As a control, lysates were also analyzed with antibodies against β-actin. (G) Intensity of the bands in panel F were determined with a Thermo Fisher iBright 1500 Imaging System and plotted as the ratio of ASP-expressing over control cells. (H) Impact of ASP on maintenance of mitochondrial membrane potential following treatment with the protonophore, CCCP. HeLa cells stably transduced with an ASP-expressing lentiviral vector (HeLa-ASP) or an empty lentiviral vector (HeLa-EV) were stained with LIVE/DEAD™ stain, treated with 5 μM CCCP or vehicle (DMSO), and then with 80 nM TMRM. Geometric mean fluorescence intensity (GMFI) in the mock- and CCCP-treated samples was determined using FlowJo after exclusion of cell doublets and dead cells. The graph shows the percent reduction of GMFI of the lipophilic fluorescent dye, TMRM as a function of the change in mitochondrial membrane potential (ΔΨm) following CCCP-induced depolarization. The assay was performed in 5 independent biological replicates. Statistical analyses were performed with GraphPad Prism 10.6.

Four lines of evidence support the hypothesis that this observation reflects a specific effect of ASP on mitochondrial protein translation rather than through other mechanisms. First, ASP does not alter mtDNA copy number, which could account for increased mtDNA-encoded protein levels. Indeed, quantitative PCR (qPCR) assays did not show any difference in the copy number of three mtDNA-encoded genes (ATP6, ND1, and CYTB) relative to that of two nDNA-encoded genes (albumin and β-globin, invariably present at 2 copies per cells) in ASP-expressing compared to control cells (Fig. S9). Second, RT-qPCR assays showed that ASP does not significantly affect the transcription levels of mtDNA-encoded ATP6, ND1, and CYTB relative to mRNA levels of nDNA-encoded GAPDH (Fig. S10). Third, western blot analyses showed that ASP does impact the expression levels of 5 subunits of mitochondrial complexes I (NDUFS1), III (RISP), IV (COX4), and V (ATP5A1), and the Complex III-IV electron carrier, cytochrome c that are encoded in the nuclear genome (Fig. 6D, E). Fourth, the expression of 3 other HIV-1 proteins (TAT, VPR, GAG) did not have an impact on the expression of mtDNA-encoded proteins ATP8, COX1, COX2, and Cyt-b expression similar to ASP (Fig. 6F, G), showing that this effect is not a cellular response to ectopic expression of an exogenous protein.

Next, we sought to test if ASP interactions affect mitochondrial function. We therefore examined mitochondrial membrane potential (ΔΨm) under basal and stress conditions using tetramethylrhodamine methyl ester (TMRM) in flow cytometry. Basal TMRM fluorescence was comparable between the two cell lines (Fig. S11A), indicating that ASP expression did not substantially alter resting mitochondrial membrane potential. However, treatment with carbonyl cyanide m-chlorophenyl hydrazone (CCCP) resulted in a significantly smaller decrease in TMRM fluorescence in ASP-expressing cells compared to control cells. At 5 μM CCCP, TMRM fluorescence decreased by approximately 55–60% in control cells, whereas the decrease was 10– 15 percentage points smaller in ASP-expressing cells (P = 0.0026, n = 5 biological replicates; Fig. 6H). A smaller (3–10 percentage-point reduction) but significant difference was also observed at 20 μM CCCP (P = 0.0041, n = 6 biological replicates; Fig. S11B). These results indicate that, although basal ΔΨm was similar between the two cell lines, ASP-expressing cells were more resistant to CCCP-induced mitochondrial depolarization. Consistent with this result, oxygen consumption rate (OCR) measurements showed a reproducible trend toward higher spare respiratory capacity (difference between FCCP-stimulated maximum and basal respiration) in ASP-expressing cells compared with control cells (Fig. S11C).

These findings are consistent with a model in which ASP-associated mitochondrial ribosome interactions alter mitochondrial respiratory-chain function and increase the capacity of mitochondria to withstand depolarizing stress.

## Discussion

Here, we report a first, comprehensive characterization of the human ASP protein interactome, uncovering functional roles for ASP in a number of related physiological processes that ensure cell homeostasis. We propose that ASP may promote viral persistence by supporting the fitness and by extending the lifespan of the infected cell.

Previous investigation into a functional role of ASP has been hindered by a lack of suitable reagents and the low expression levels of ASP *in vivo* and *in vitro* due to weak transcriptional activity from the negative sense promoter in the proviral HIV-1 3’ LTR (*6, 35*). In this study, we have overcome the lack of suitable antibodies for co-immunoprecipitation studies by creating a C-terminal EPEA tag in the *asp* ORF of the JRCSF molecular clone. This allowed us to identify ASP binding partners by expressing ASP under its natural promoter in the HIV-1JRCSF molecular clone. For functional validations, we took advantage of robust, ectopic ASP overexpression. Using this approach, we demonstrate that ASP has the capacity to protect from DNA damage and stabilize mitochondrial membrane potential in response to acute experimental perturbations. We note that ASP expression and its physiological effects are expected to be more subtle in HIV-infected cells, reflecting lower levels of chronic stress. Nevertheless, their cumulative impact may become increasingly significant over the prolonged course of HIV-1 infection (*7*).

Our previous work showed that knocking out the expression of ASP reduces the rate of cell-free viral entry (*15*). This is consistent with the evidence that ASP is present on the envelope of infectious virions (*7*) and suggests that, while ASP is not required for viral entry, it enhances this early step of the replication cycle. In the present study, we have focused our efforts on potential role(s) of intracellular ASP. The picture that emerges from this analysis is that ASP interacts with functional pathways related to cellular stress responses and cell homeostasis, including chromatin maintenance and DNA repair, mitochondrial protein translation, and RNA metabolism and processing. Given that ASP interacts with both DNA repair factors and repair-relevant chromatin remodelers, we speculate that ASP may help establish a repair-conducive chromatin environment and/or modulate the activity and/or retention of repair factors at sites of damage. However, we cannot formally rule out an indirect role for ASP, for example through altered mRNA processing of repair factors. The information provided by our proteomic screen will be instrumental in guiding future efforts to elucidate how ASP promotes genome integrity. It will further be interesting to determine if ASP affects chromatin transactions other than DNA repair.

Consistent with its nuclear functions, our previous study showed that ASP localizes exclusively to the nucleus during viral latency. However, it is also found in the cytoplasm following viral restimulation. Consistent with active transport between nucleus and cytoplasm, we show that the karyopherin TNPO1 interacts with and contributes to the nuclear import of ASP. What controls ASP shuttling between the two compartments, and which other import pathways may contribute to account for the residual nuclear localization of ASP observed in the absence of TNPO1 remains to be determined (*36*).

The functional modules identified in our study are not disconnected from each other. Indeed, several proteins in our dataset are known to integrate and bridge the functional pathways. Prominent examples are the three subunits of the RPA complex (RPA70, RPA32, and RPA14), which has been shown to link DNA damage and transcription stress (*37, 38*). In addition, the two subunits of the FACT complex (SSRP1 and SUPT16H) impact both transcription and DNA repair through chromatin remodeling (*39*). Other examples include DHX36, UPF1, WDR5, RBBP5, and DPY30, which link chromatin, genome stability, RNA surveillance, and RNA processing (*37, 40–43*). These findings provide a framework for the long overdue process of unraveling the function of an HIV-1 protein that remains to be fully characterized.

The diverse cellular distribution and pleiotropic activities of ASP that emerge from this and our previous study are not unique. Other proteins of HIV-1 show similar behavior. A few examples include Tat, which shuttles between nucleus, nucleolus, cytoplasm, and extracellular space, displaying both intracellular and extracellular activities (*44*), Vpr, which has functions in the nucleus, nuclear envelope, cytoplasm, mitochondria, chromatin-associated complexes, and virions (*45*), and Nef, which localizes to the plasma membrane, cytoplasm, endosomes, Golgi-associated vesicles, and endocytic trafficking compartments (*46*). This functional pleiotropy extends beyond HIV-1 to viral proteins such as HPV E6 and E7 and EBV EBNA, which have been shown to impact multiple cellular functional pathways by interacting with a broad array of cellular proteins. These viral “multitasking” proteins often contain intrinsically disordered regions (*47*) and short linear interaction motifs (*48, 49*) that enable flexible binding to diverse host proteins involved in immunity, transcription, trafficking, metabolism, and cell-cycle regulation. The presence of short predicted intrinsically disordered regions within ASP (*50*) may facilitate its interactions with diverse cellular proteins.

Overall, the studies described here corroborate the functional relevance of ASP and are expected to open new avenues of investigation on a viral protein that has remained largely uncharacterized, fostering a deeper understanding of the mechanisms through which HIV-1 hijacks cellular functions and achieves persistent infection.

## Author contributions

Conceptualization: MSI, THL, PO, FR

Methodology: MSI, THL, CLL, RL, XL, IC, PO, FR

Investigation: MSI, THL, CLL, XJ, YW, PO, FR

Visualization: THL, CLL, PO, FR

Funding acquisition: PO, FR

Project administration: PO, FR

Supervision: PO, FR

Writing – original draft: PO, FR

Writing – review & editing: MSI, THL, CLL, YW, XJ, RL, XL, IC, PO, FR

## Data, code, and materials availability

All data are available in the main text or the supplementary materials. All material generated in this study (including but not limited to oligonucleotides, plasmids and cell lines) are available on request.

## Acknowledgments

We acknowledge the use of artificial intelligence technology to fit the main text within the word limit allowed by the journal.

## Supplementary Materials

Materials and Methods

Figs. S1 to S11

Tables S1 and S2

Data S1

References (51–56)

