## Supplementary Materials for "A proteomics screen of HIV-1 ASP interactors reveals multifaceted effects on host cell homeostasis"

**The PDF file includes:**

Materials and Methods
Supplementary Text
Figs. S1 to S11
Tables S1 and S2
References *51* to *56*

**Other Supplementary Materials for this manuscript include the following:**

Data S1

### Materials and Methods

**Cell lines.** HEK293T and HeLa cells were obtained from American Type Culture Collection. Cells were cultured at 37°C in 5% $CO_2$ atmosphere in complete DMEM medium containing 90% Dulbecco's Modified Eagle Medium (DMEM) and 10% heat-inactivated fetal bovine serum (FBS), supplemented with 100 units/ ml penicillin, 100 mg/ml streptomycin, and 2 mM L-Glutamine (all from ThermoFisher). Cells were cultured for up to 6-8 passages and then discarded. Frozen cell stocks were prepared in 90% FBS and 10% dimethyl sulfoxide (DMSO, ThermoFisher) and stored at –182°C in liquid nitrogen.

**Plasmids.** To generate the HIV-1 molecular clone expressing EPEA-tagged ASP, the pYK-JRCSF vector (ARP-2708, HIV Reagent Program) (*17*, *51*) was digested with *Bam*HI and *Ale*I to excise a 700-bp fragment within the HIV-1 genome that encompasses the ASP open reading frame. Then, a synthetic 709-bp fragment flanked by *Bam*HI and *Ale*I restriction sites was inserted into the linearized vector. The sequence of the 709-bp fragment is identical as the 700-bp excised fragment except for a new 9-bp sequence that replaces nucleotides 5'-CTGTAA-3' at positions 1140-1145 of *env* in the original vector with 5'-CGCTTCTGG-3'. This substitution changed codons 182 and 183 in the *asp* ORF from 5'-TTA CAG-3' to 5'-CCA GAA-3' (L182P Q183E) and inserted a new GCG codon (184A) before the TAG *stop* codon. Altogether, the new 9-bp sequence inserted an EPEA tag (glutamic acid – proline – glutamic acid – alanine) at the C-terminus of ASP, while leaving the rest of the HIV-1 genomic sequence unaffected. The sequence was verified via Sanger sequencing with forward primer 5'-CATTAGTAGAGCACAATGGAATAACAC-3'; reverse primer 5'-CCATCCAGGTCATGTTATTCCAAATAC-3'. The eukaryotic vector expressing ASP with three FLAG epitopes (DYKDHDG-DYKDHDI-DYKDDDDK) appended at the C-terminus was purchased from VectorBuilder. This pCAG-ASP-3×FLAG vector expresses ASP-3×FLAG under the CAG promoter (CMV early enhancer, chicken β-globin promoter, rabbit b-globin splice acceptor) and Enhanced Green Fluorescent Protein (EGFP) under the human cytomegalovirus immediate early enhancer/promoter. The ASP-DPY-NLS vector was derived from pCAG-ASP-3×FLAG. The vector was digested with *Bbv*CI and *Cla*I to excise a 559-bp fragment encompassing the ASP open reading frame. Then a synthetic fragment of equal length and flanked by the same restriction sites was inserted. This fragment created mutations in three proline-tyrosine nuclear localization signals (PY-NLS) motifs in the ASP sequence. PY-NLS1 (aa 41-59): P43A, I44A, R54A, P58A, I59A; PY-NLS2 (aa 96-111): L96A, S97A, L98A, I99A, K106A, P110A, L111A; PY-NLS3 (aa 116-134): P116A, S117A, L118A, V119A, H129A, P133A, L134A. The pLV-ASP-3×FLAG vector carrying the puromycin resistance gene was purchased from VectorBuilder.

**Transfection and transduction.** The day before transfection, $4 \times 10^6$ HEK293T cells were plated on 10-cm tissue culture dishes in 10 ml of complete DMEM medium to achieve ~70% confluency the following day. The day of the transfection, the complete DMEM medium was removed and replaced with 10 ml of serum-free Opti-MEM. For each plate, 12 mg of plasmid DNA were diluted in 1.5 ml of Opti-MEM and 36 mg of PEI-Max were diluted in 1.5 ml of Opti-MEM in a separate tube. The two tubes were mixed together and incubated at room temperature for 15 minutes. Then, the 3-ml mix was added dropwise onto the cells. After 6-8 hours, the Opti-MEM medium was replaced with complete DMEM medium, and the cells were analyzed 48 hours later. For lentiviral packaging, cells were transfected with a mixture of transfer vector (pLV-ASP-3×FLAG), packaging vector (psPAX2; Addgene plasmid #12260), and pseudotyping vector (VSV-G) at 4:3:1 mass ratio (12 mg total DNA amount). For purification of lentiviral particles, culture medium was harvested 48 hours post-transfection and centrifuged at 2000 rpm for 15 minutes at 4°C to remove cell debris. The centrifugation supernatant was then passed through a 0.2-mm filter, mixed with 0.2 volumes PEG-it Virus Precipitation Solution (5×; System Biosciences), and incubated overnight at 4°C. The next day, the solution was centrifuged at 1500 × *g* for 30 minutes at 4°C. The viral pellet was resuspended in 1/100$^{th}$ volume of sterile 1× PBS and used immediately or stored at -80°C. For transduction, 1 × $10^6$ HeLa cells were mixed with the concentrated lentiviral preparation, spinoculated at 1,200 × *g* for 30 minutes at room temperature and then cultured in complete DMEM medium. After 48 hours, the culture medium was supplemented with 1 mg/ml puromycin to select stably transduced cells, which were thereafter maintained in selection medium.

**ASP-EPEA capture and mass spectrometry.** Cells transfected with the $JRCSF_{WT}$ and the $JRCSF_{ASP\text{-}EPEA}$ vectors were harvested 48 hours after transfection and lysed with Pierce IP Lysis Buffer under non-denaturing conditions and in the presence of protease inhibitors. Next, equal amounts of cell lysates were incubated with CaptureSelect™ C-tagXL Affinity Matrix (anti-EPEA; ThermoFisher) to capture and elute EPEA-tagged ASP according to the manufacturer's instructions. Lysates from cells transfected with the $JRCSF_{WT}$ vector served as background controls. Proteins eluted from C-tagXL affinity matrix were analyzed by mass spectrometry as described previously with modifications (*52*, *53*). Protein samples (50 mg) were brought up to 100 ml with 10 mM TEAB plus 1% SDS and disulfide bonds reduced with 10 ml of 50 mM dithiothreitol for 1 hour at 60°C. Samples were cooled to RT and pH adjusted to ~7.5, followed by alkylation with 10 ml of 100 mM iodoacetamide in the dark at room temperature for 15 minutes. Proteins were then precipitated with TCA/acetone at -20°C overnight. The pellets were washed with acetone, reconstituted in 3 ml of 500 mM TEAB and 15 ml of water, sonicated 10 for minutes, and proteolyzed with 800 ng of MS grade trypsin (Thermo-Pierce) in 20 ml of 75mM TEAB at 37°C overnight (*54*). Peptides were acidified with 0.1% TFA, desalted on Oasis HLB mElution 96-well plates, eluted with 50 ml of 60% ACN/0.1% TFA, and dried. Desalted peptides were re-constituted in 20 ml of 2% acetonitrile/0.1% formic acid and diluted to 250 ng/ml. Peptides were analyzed on nano-Orbitrap Fusion Lumos-ETD in FTFT (Thermo Fisher Scientific) interfaced with nano-LC 1200 system (ThermoFisher) using reverse-phase chromatography at 300 nl/min, on 75 mm × 150 mm ProntoSIL-120-5-C18 H column 3 mm, 120Å (BISCHOFF). Eluted peptides were sprayed into an Lumos-ETD mass spectrometer through 1 mm emitter tip (New Objective) at 2.4 kV. Survey scans (Full MS) were acquired on Orbi-trap within 360-1700Da m/z using Data-Dependent Aquisition "Top 15" method with dynamic exclusion of 15 s. Precursor ions were individually isolated with 0.8 Da (no offset) and fragmented (MS/MS) using an HCD activation with collision energy 30. Tandem MS2 spectra were processed by Proteome Discoverer (v2.4 ThermoFisher Scientific) using fileRC option (re-calibration) and analyzed with Mascot v.2.7 Matrix Science using Target Decoy Validator for PSM validation. Mascot database search was done against RefSeq2021_204_Human. Mascot .dat files were sent to PD2.4 for PSM validation with TDV and then loaded on Scaffold.

**Co-immunoprecipitation and Western blot.** Co-IP assays were carried out with 500 μg of whole cell lysate diluted to 500 μl with PBS. Ten micrograms of target-specific antibody or isotype control (Table S1) were added to each reaction, followed by 4-hour incubation at 4°C. Subsequently, 100 μl of Protein A/G Dynabeads (Thermo Fisher) were added to each sample, followed by an overnight incubation at 4°C with end-over-end rotation. Immunocomplexes were then immobilized with a magnetic rack and washed three times with TBS-T buffer (20 mM Tris base, 150 mM NaCl, 0.1% Tween-20) supplemented with 1 mM EDTA and protease inhibitor cocktail (Roche Biochemicals). Immunocomplexes were then eluted by adding SDS-PAGE loading buffer followed by incubation at 37°C for 1 hour or at 70°C for 15 minutes. Co-IP samples (along with 10% of whole cell lysate as Input) were run on a 4-20% Tris-Glycine polyacrylamide gel, transferred to a PVDF membrane (90 minutes at 60 V), probed with primary and secondary antibodies (Table S1), and analyzed by ECL (Pierce). Images were captured and analyzed using an iBright CL1500 Imaging System (Thermo Fisher).

**Alkaline comet assay.** The alkaline comet assay was conducted following the manufacturer's instruction for the Trevigen CometAssay™ kit. Briefly, cells were harvested by trypsinization at 37°C for 2 minutes following the indicated treatment and release conditions. After centrifugation, the cell pellet was resuspended in ice-cold PBS to a density of $5 \times 10^5$ cells/mL and mixed with LMAgarose (R&D systems) at 37°C at a ratio of 1:10 (v/v). This mixture was immediately transferred onto CometSlides (R&D systems), allowed to solidify, and immersed in prechilled lysis solution (R&D systems) overnight at 4°C. For DNA denaturation, the slides were incubated in alkaline unwinding solution (200 mM NaOH, 1 mM EDTA) for 20 minutes at RT, followed by electrophoresis (25V for 30 minutes) in alkaline electrophoresis buffer (200 mM NaOH, 1 mM EDTA) at 4°C. Following electrophoresis, the slides were rinsed twice with distilled $H_2O$, once with 70% EtOH, and air-dried before being stained with SYBR® Gold in TE buffer for 30 minutes. Finally, the slides were mounted with ProLong Gold Antifade Mountant (ThermoFisher) and visualized under a Zeiss Axio Imager Z1 microscope (acquisition software ZEN3.7). Statistical analysis was performed using OpenComet plugin in ImageJ.

**Immunofluorescence.** For immunofluorescence (IF) of 8-oxoG (*55*), HeLa cells were plated on poly-L-lysine-coated coverslips 24 hours prior to treatment with 500 μM $H_2O_2$ for 30 minutes, followed by a 0 or 4-hour release time. Cells were fixed in an acetone:methanol (1:1) mixture, air-dried, and rehydrated in 1× PBS for 15 minutes. DNA denaturation was performed by incubating the cells in 2N HCl for 45 minutes at RT. After washing three times with 1× PBS, cells were neutralized with 50 mM Tris-HCl (pH 8.8) for 5 minutes. Following neutralization, cells were permeabilized with 0.5% Triton X-100 in 1× PBS for 10 minutes, washed with 1× PBS-T (0.1% Triton X-100) and blocked with 20% FBS in 1× PBS-T for 1 hour at RT. After blocking, cells were washed and incubated with a mouse anti-8-oxoG antibody (Santa Cruz, sc-130914) at 4°C overnight. The following day, cells were washed with 1× PBS-T and incubated with an Alexa 594-conjugated anti-mouse secondary antibody (Invitrogen, A-11005) diluted in 1× PBS-T, 5% FBS for 1 hour at RT. Coverslips were immersed in 1× PBS with 5 μg/mL of Hoechst 33342 (Sigma-Aldrich) for 5 minutes and mounted on slides with ProLong Gold Antifade Mountant (ThermoFisher). Images were captured with Zeiss LSM 900 with Airyscan2 (acquisition software ZEN3.6). For 8-oxoG intensity analysis, 100-150 nuclei were randomly selected per condition, and their nuclear intensities were quantified using ImageJ.

**Subcellular fractionation.** For nuclear and chromatin fractionation, $1 \times 10^7$ cells were lysed in ice-cold cytoplasmic buffer (10 mM HEPES-KOH pH 7.6, 3 mM $MgCl_2$, 40 mM KCl, 2 mM DTT, 5% glycerol, 0.5% NP-40, cOmplete™ protease inhibitor cocktail (Roche)) for 10 minutes on ice, followed by centrifugation at $1,250 \times g$ for 5 minutes at 4°C. The pellet was washed once with the cytoplasmic buffer, centrifuged again and resuspended in nuclear lysis buffer (10 mM HEPES-KOH pH 7.9, 0.1 mM EGTA, 1.5 mM $MgCl_2$, 420 mM NaCl, 0.5 mM DTT, 25% glycerol, cOmplete™ protease inhibitor cocktail). After incubation on ice for 10 minutes, the nuclear lysates were centrifuged at $1,250 \times g$ for 5 minutes at 4°C. The supernatant was collected as the nuclear fraction. The remaining chromatin-containing pellet was then resuspended in N-buffer (20 mM Tris-HCl pH 7.5, 1 mM $CaCl_2$, 100 mM KCl, 0.3 M sucrose, 2 mM $MgCl_2$, 0.1% Triton X-100, and cOmplete™ protease inhibitor cocktail) supplemented with a DNA-shearing enzyme cocktail: DNase I (10 U/µL, Promega), micrococcal nuclease (40 U/µL, NEB), RNase A (20 ng, Qiagen), and BSA (1:100, Promega). Chromatin was solubilized by pipetting and incubated at 30°C for 30 minutes. Finally, the lysates were centrifuged at $13,000 \times g$ for 10 minutes at RT, and the supernatant was collected as the chromatin fraction. Protein concentrations were determined using the Bradford assay.

**Cell viability assays.** For MTT assay, cells were seeded in 96-well plates at $3 \times 10^3$ cells/well. For $H_2O_2$ or MMS treatments, cells were exposed to the indicated doses for 24 hours, washed out with 1× PBS, and incubated in fresh medium for an additional 24 hours. For UV irradiation, cells were washed with 1× PBS, aspirated, and then irradiated with the indicated UV doses, and incubated for 48 hours. To measure cell viability, the culture medium was replaced with 0.5 mg/mL MTT reagent (Sigma Aldrich), and cells were incubated for 2 hours at 37 °C. The formazan crystals were then dissolved in 100 µL of DMSO for at least 20 minutes. Absorbance was measured at 570 nm using SpectraMax M5 microplate reader (Molecular Devices), with SoftMax Pro 5.2 software (Molecular Devices).

**Extraction of DNA from MNase-digested nuclear fractions.** The nucleosome profiling assay was conducted based on a mononucleosomes co-IP protocol previously described (*56*). HeLa cells expressing either empty vector (EV) or FLAG-ASP were harvested by trypsinization and counted to obtain $8 \times 10^6$ cells for an experimental set. Cells were pelleted by centrifugation at $300 \times g$ for 5 minutes at 4°C, washed, and resuspended in ice-cold 1× PBS. Cells were then centrifuged again and lysed in 1 ml of buffer A (10 mM HEPES (pH 8.0), 10 mM KCl, 1.5 mM $MgCl_2$, 340 mM Sucrose, 10% Glycerol, 1 mM DTT, 0.2% Triton X-100) containing protease inhibitor cocktail, followed by a 5-minute incubation on ice to isolate the nuclei. The intact nuclei were centrifuged at $600 \times g$ 5 min at 4°C, washed with buffer A, and then equally aliquoted for eight distinct conditions MNase treatement conditions: four MNase concentrations (0, 0.025, 0.1, 0.4 U) across two incubation time point (5 and 20 min). MNase digestion was carried out in 500 µL of cutting buffer (15 mM NaCl, 60 mM KCl, 10 mM Tris (pH 7.5), 2 mM $CaCl_2$) at 37°C for the indicated times and terminated by the addition of 20 mM EGTA. The digested nuclei were centrifuged at 1300 × *g* for 5 minutes at 4°C to collect the initial supernatant (S1 fraction). To recover the remaining nucleosomes bound to the nuclear pellet, the pellet was subsequently incubated with 500 µL of TE buffer on ice for 30 minutes and centrifuged at 13,000 × *g* for 5 minutes at 4°C to obtain the second supernatant (S2 fraction), 100 µL aliquots of the S1 and S2 fractions from each condition were combined in a new tube. For DNA extraction, the combined fractions were treated with 1 µL of 10 mg/mL proteinase K and incubated at 37°C overnight, followed by phenol:choloform extraction and ethanol precipitationPurified DNA was loaded and run on a 1.2% agarose gel to visualize the nucleosomal DNA ladder. To quantify chromatin accessibility and the distribution of mono-, di-, tri-, tetra-, and poly-nucleosomes, densitometric profiling was performed using ImageJ. The line profile tool was utilized to measure the signal intensity (in arbitrary units, a.u.) along the migration distance (in inches) for each lane to compare the digestion patterns between the EV and FLAG-ASP groups.

**Reverse Transcription.** Reverse transcription was performed using the iScript™ Select cDNA Synthesis Kit (Bio-Rad, No. 1708897). Each reaction contained 250 ng RNA, 1 µL random primer mix, 0.5 µL reverse transcriptase enzyme, and molecular biology-grade water to a final volume of 10 µL. Negative control reactions were prepared without reverse transcriptase enzyme to confirm the absence of genomic DNA contamination. Reverse transcription was then performed under the following conditions: 25°C for 5 minutes, 42°C for 30 minutes, 85°C for 5 minutes, and 4°C to hold.

**Quantitative PCR.** Quantitative PCR reactions were performed using Taq Pro Universal SYBR qPCR Master Mix (Vazyme, No. Q712-02). Each reaction contained: 5 µL Taq Pro Universal SYBR qPCR Master Mix, 0.5 µL forward and reverse primer mix (Table S2), 3.5 µL molecular biology-grade water, and 1 µL template DNA or cDNA. Negative control reactions contained water instead of the template. In quantitative PCR, cDNA samples were denatured at 95°C for 2 minutes. Amplification was performed for 35 cycles, each consisting of denaturation at 95°C for 10 seconds and annealing/extension at 60°C for 40 seconds. After amplification, a melting curve was obtained by holding the samples at 95°C for 15 seconds, cooling to 60°C for 60 seconds, and ramping to 95°C for 15 seconds while fluorescence was recorded. Threshold cycle (Ct) values were obtained from quantitative PCR, and relative fold changes were calculated using the comparative $2^{-\Delta\Delta Ct}$ method.

**Mitochondrial function assays.** Mitochondrial respiration was assessed using the Extracellular Oxygen Consumption Assay Kit (Cayman Chemical, No. 502880) according to the manufacturer's protocol, with the modification that the volume of oxygen probe added per well was reduced from 5 µL to 2 µL, as 2 µL was sufficient to generate a strong fluorescence signal clearly distinguishable from baseline. The pharmacological inhibitors used in these assays were: carbonyl cyanide-p-trifluoromethoxy phenylhydrazone (FCCP; Cayman Chemical, No. 15218), Oligomycin (Cayman Chemical, No. 11341), and Antimycin A (Cayman Chemical, No. 34799). All three were reconstituted in dimethyl sulfoxide (DMSO; Sigma-Aldrich, No. D2650). All treatments, including the vehicle (DMSO) control, contained a final concentration of 0.5% DMSO to ensure consistency across conditions. HeLa-ASP and HeLa-EV cells were seeded at a density of 60,000 cells per well in a 96-well plate and incubated for 24 hours at 37°C with 5% $CO_2$ in 200 µL complete Dulbecco's Modified Eagle Medium (cDMEM) consisting of DMEM (Gibco™, No. 11965092) supplemented with 10% heat-inactivated fetal bovine serum (Bio-Techne®, No. S11150), 1% penicillin-streptomycin (Gibco™, No. 15140122) to allow cell attachment and recovery prior to assay. After 24 hours of incubation, treatment solutions were prepared in complete media such that final in-well concentrations were 1.5 µM oligomycin, 1.0 µM FCCP, 1.0 µM antimycin A, and 0.5% DMSO for all conditions, including vehicle. For each well, a master mix containing 10 µL drug solution, 2 µL oxygen probe, and 88 µL media was prepared. The optimal concentration of FCCP was determined via a titration experiment ranging from 0.1 µM to 5 µM with wild-type HeLa cells. A final 1.0 µM FCCP concentration yielded the steepest slope of the fluorescence curve during which oxygen consumption increases. Culture media was aspirated from each well, and 100 µL of the prepared mixture was added. Each well was then sealed with approximately 100 µL of HS mineral oil prewarmed to 37°C. The plate was immediately transferred to the reader chamber, and fluorescence was measured for 2 hours at 5-minute intervals under previously optimized conditions using a SpectraMax® iD5 plate reader (Molecular Devices) with SoftMax Pro 7.1.2 software. Signal optimization experiments demonstrated that the oxygen probe produced a detectable signal that was clearly separated from blank wells under the following settings: fluorescent kinetic mode, 37°C, excitation at 380 nm, emission at 650 nm, medium photomultiplier tube (PMT) gain, top read with a read height of 5.0 mm, and an integration time of 400 ms. These parameters were selected to maximize signal-to-noise ratio while minimizing signal saturation and variability. Spare respiratory capacity was calculated as the difference between maximum (FCCP-stimulated) OCR and basal (vehicle-treated) OCR. Since these parameters were calculated as the difference between two treatment conditions, and since the number of technical replicates differed between treatment groups, conventional calculations of standard deviation followed by a t-test were not appropriate. Thus, we applied error propagation to estimate the standard error of derived parameters, followed by statistical comparison using Welch's unequal-variance t-test. Analysis of mitochondrial membrane potential state was performed using the MitoProbe™ TMRM Assay Kit for Flow Cytometry (Thermo Fisher) following the manufacturer's recommendations with minor modifications. For each sample, $1\times10^6$ HeLa-ASP or HeLa-EV cells were first stained with the LIVE/DEAD™ Fixable Aqua Dead Cell Stain Kit (Thermo Fisher). After centrifugation at 200 × *g* for 5 minutes at room temperature, cells were resuspended in 1 ml of PBS and treated with 5 µM or 20 µM carbonyl cyanide 3-chlorophenylhydrazone (CCCP) or vehicle (DMSO) for 5 minutes at 37°C, 5% $CO_2$. Subsequently, tetramethylrhodamine methyl ester (TMRM) was added at a final concentration of 80 nM, and the cells were incubated for 30 minutes at 37°C, 5% $CO_2$. Finally, cells were centrifuged at 200 × *g* for 5 minutes at room temperature, washed with 1 ml of PBS, resuspended in 0.5 ml of PBS, and immediately analyzed by flow cytometry on a LSRII Fortessa flow cytometer (BD Biosciences, San Jose, CA). Raw data were analyzed with FlowJo software. Doublet cells were excluded using forward scatter and side scatter, and dead cells were excluded using the LIVE/DEAD™ Stain.

**Statistical analyses.** Mass spectrometry data from the two biological replicate experiments were analyzed using a two-way statistical model incorporating experimental date and bait condition. Protein enrichment in the tagged-bait pull-down was assessed relative to the corresponding untagged-bait control, with fold change, p-value, and 95% confidence intervals calculated for each protein. Proteins exhibiting ≥2-fold enrichment with a p-value <0.05 were considered significantly enriched. Proteins meeting these criteria at both input amounts and in both biological replicates were classified as high-confidence ("true") interactors. Protein–protein interaction (PPI) enrichment was assessed using STRING. The observed number of interactions among the input proteins was compared with the number of interactions expected by chance based on the connectivity (node-degree distribution) of the background protein interaction network. Statistical significance was assessed using the STRING PPI-enrichment P-value. All other biostatistical analyses were performed with GraphPad Prism (v10.6).

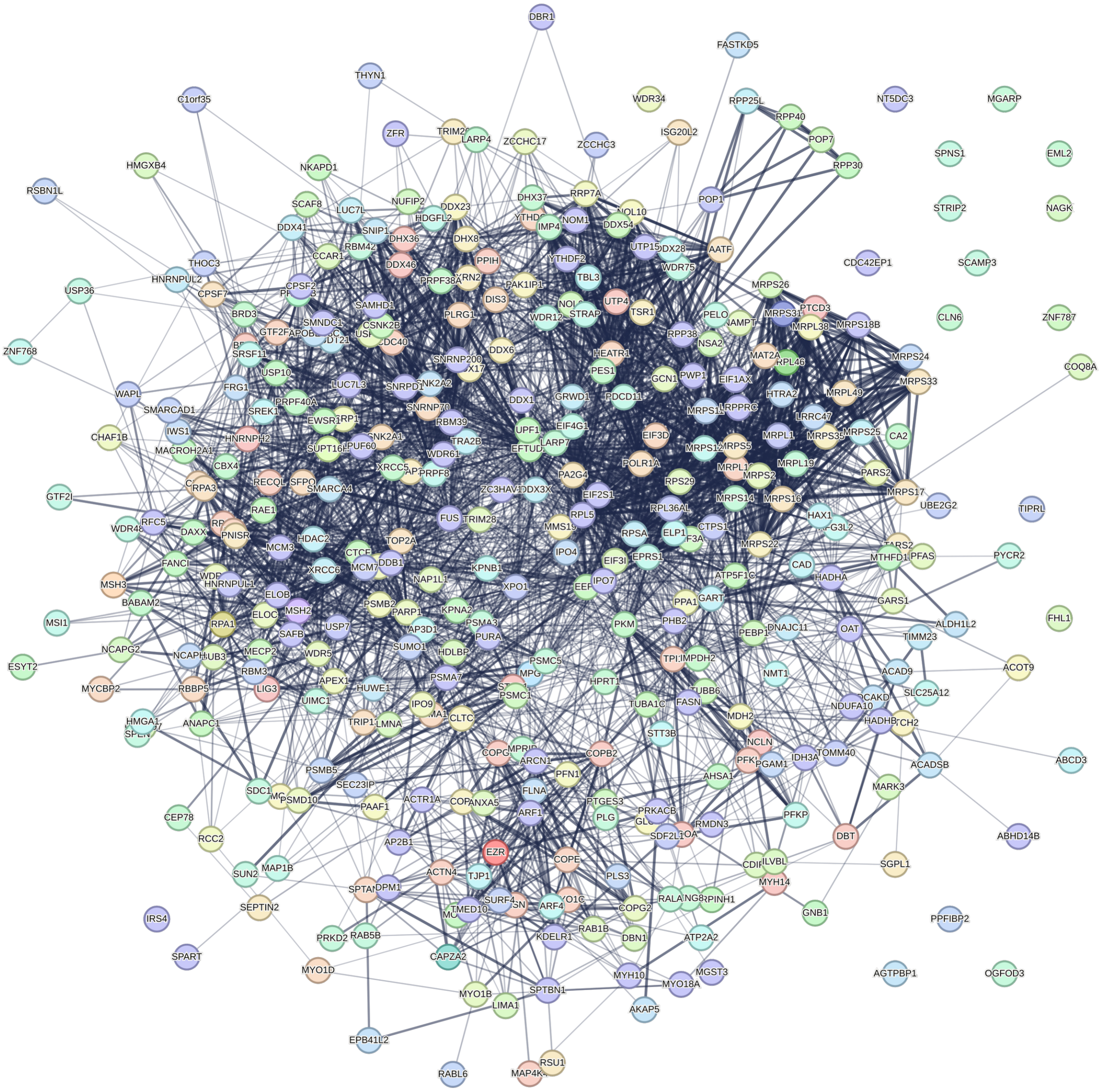


**Fig. S1.**

STRING network of genes enriched in ASP interactome. 376 host proteins are significantly enriched (≥ 2-fold, $p < 0.05$) in CaptureSelect™-enriched fractions from $JRCSF_{ASP\text{-}EPEA}$- vs $JRCSF_{WT}$-transfected 293T cells in 2 biological replicates. ASP interactors were identified by mass spectrometry of peptides obtained following trypsin digestion of CaptureSelect™-enriched proteins..

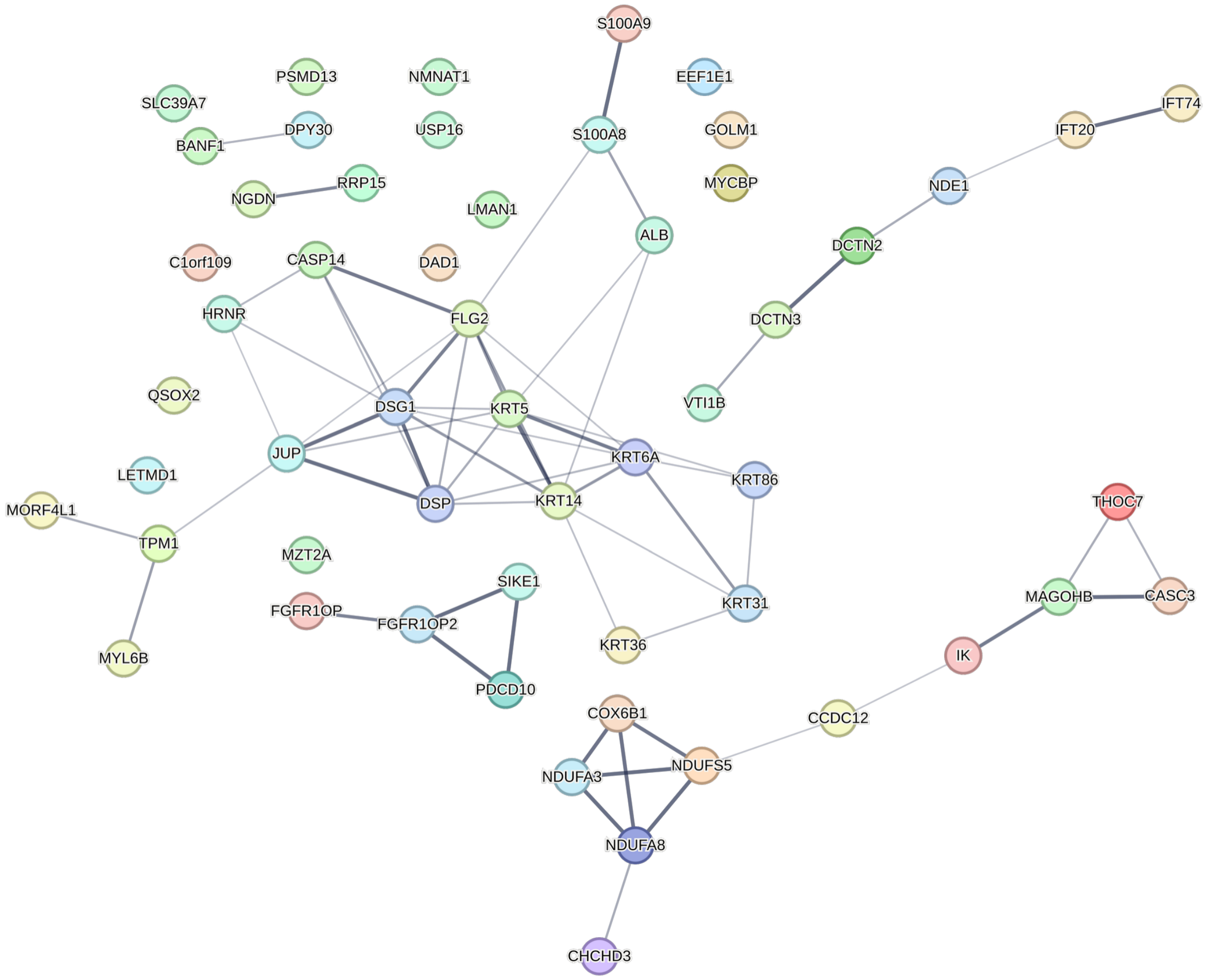


**Fig. S2.**

STRING network genes depleted in ASP interactome. 55 host proteins were significantly enriched (≥ 2-fold, $p < 0.05$) in CaptureSelect™-enriched fractions from JRCSF$_{WT}$- vs. JRCSF$_{ASP\text{-}EPEA}$-transfected 293T cells in 2 biological replicates. ASP interactors were identified by mass spectrometry of peptides obtained following trypsin digestion of CaptureSelect™-enriched proteins.

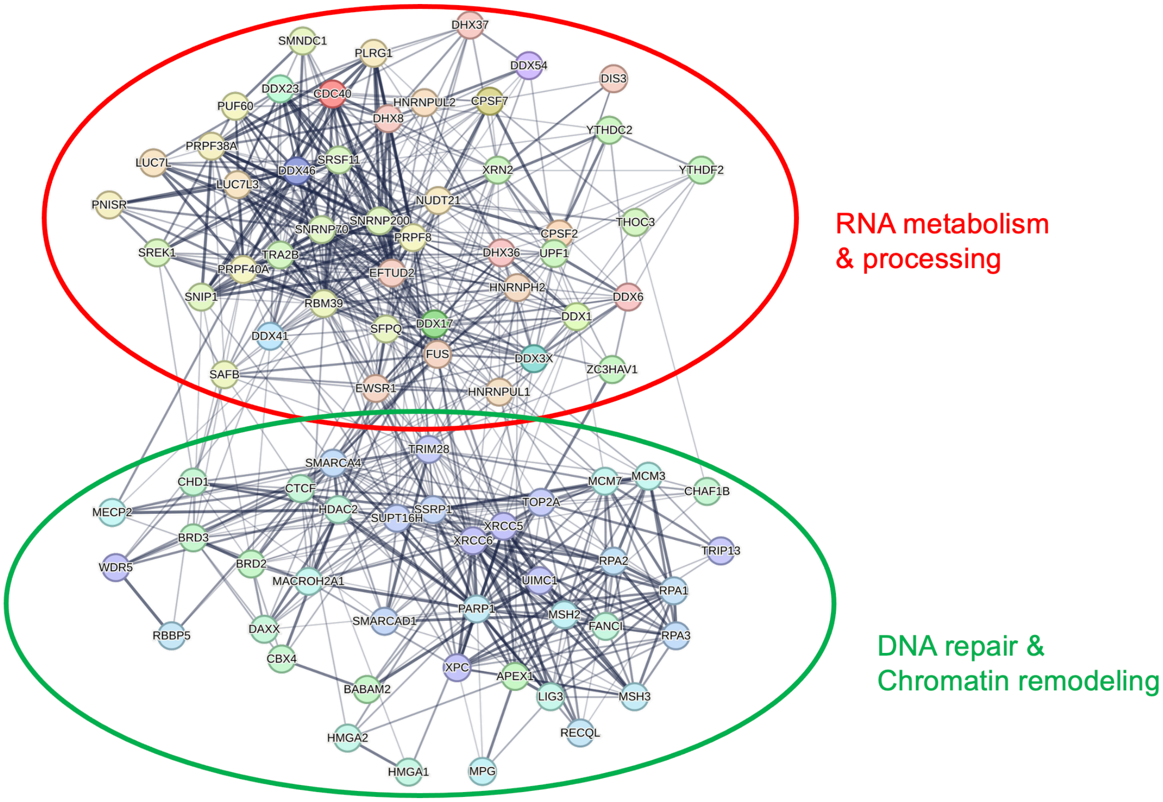


**Fig. S3.**

ASP interactome is enriched for genes involved in RNA and DNA metabolism. STRING network of 87 (~25% of total) host proteins significantly enriched (≥ 2-fold, $p < 0.05$) in CaptureSelect™-enriched fractions from $JRCSF_{ASP\text{-}EPEA}$- vs $JRCSF_{WT}$-transfected 293T cells that belong to two major nuclear functional pathways: RNA metabolism (red oval) and DNA repair & chromatin remodeling (green oval).

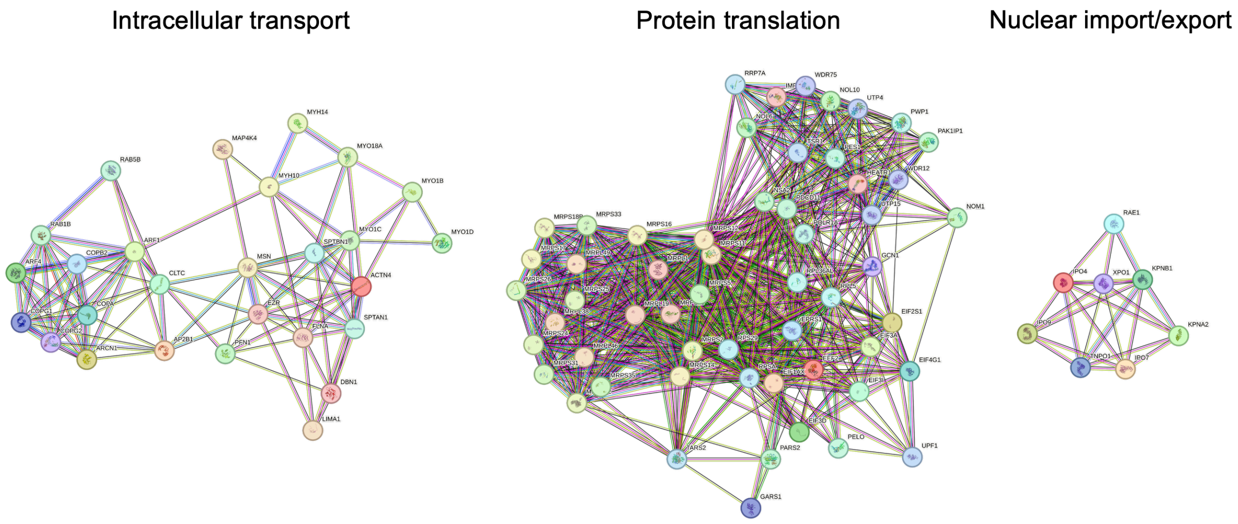


**Fig. S4.**

Additional ASP interactome pathways. STRING network of host proteins significantly enriched ($\geq$ 2-fold, $p < 0.05$) in CaptureSelect™-enriched fractions from JRCSF$_{ASP\text{-}EPEA}$- vs JRCSF$_{WT}$-transfected 293T cells that belong to three cytosolic/mitochondrial functional pathways: intracellular transport, cytosolic and mitochondrial protein translation, nuclear protein import/export.

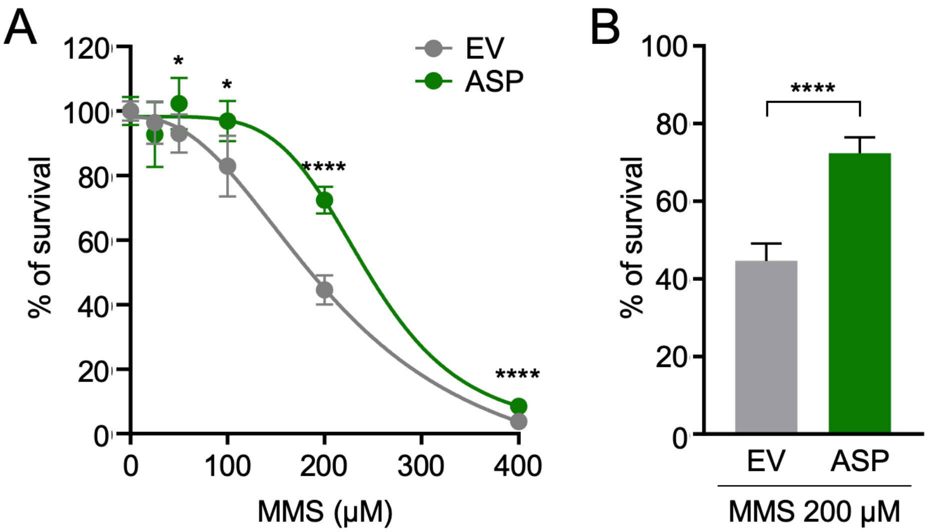


**Fig. S5.**

ASP protects from MMS-induced DNA damage. (A) Cell viability assay (MTT) in HeLa cells as in Fig. 3F. Cells were exposed to the indicated doses of MMS for 24 h, followed by 24 h recovery. Values are normalized to no treatment and expressed as mean and SD ($n = 4$ technical replicates). (B) Bar chart depicts values at 200 µM MMS. P values are based on Student's two-tailed t-test, * $P < 0.05$, **** $P < 0.0001$.

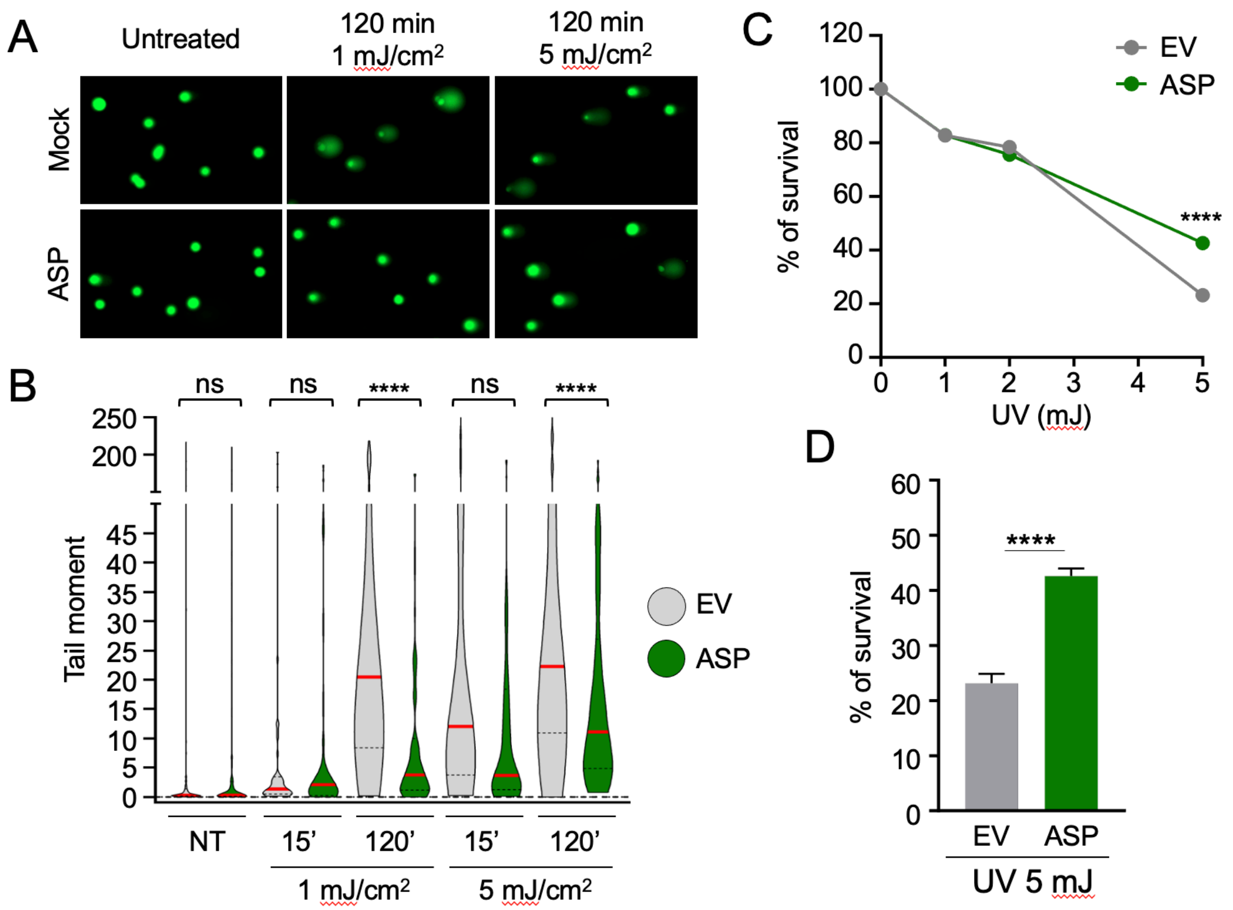


**Fig. S6.**

ASP protects from UV damage. (A) Alkaline Comet assay in HeLa cells in HeLa cells stably transduced with empty vector (EV) or ASP-3xFLAG 120 min after UV irradiation at the indicated doses, or untreated. (B) Violin plots depicting Olive tail moment, red lines indicated the median (n > 100 cells). P values are based on two-sided Mann-Whitney U test. (C) Cell viability assay (MTT) in HeLa cells as in (A) exposed to the indicated doses of UV, followed by 48 h recovery. Values are normalized to no treatment and expressed as mean and SD (n = 4 technical replicates). (D) Bar chart depicting survival fractions at 5 mJ UV. P values are based on Student’s two-tailed t-test, **** P < 0.0001.

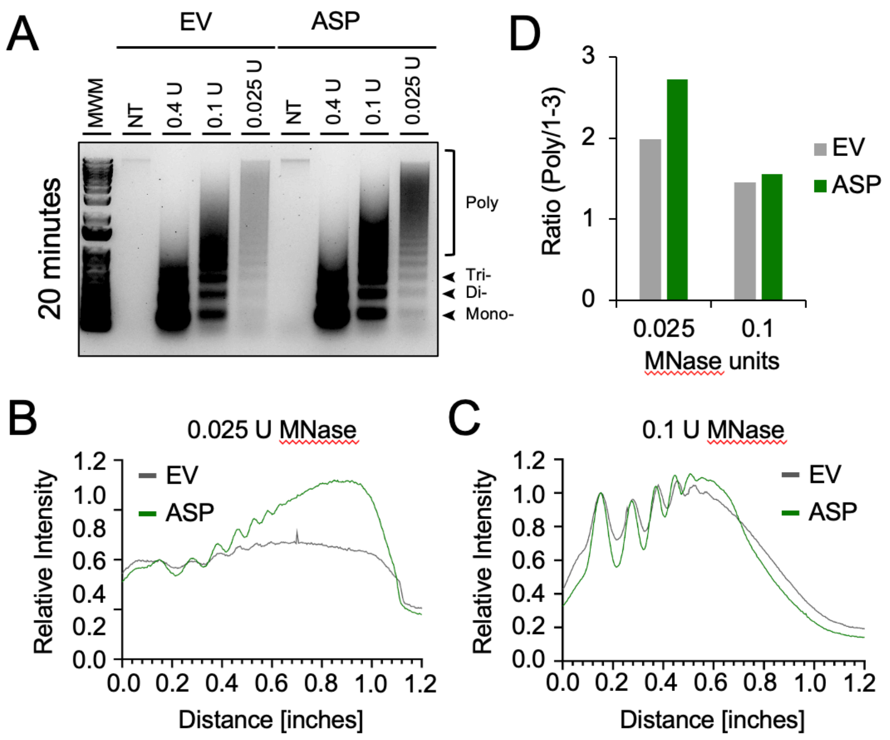


**Fig. S7.**
ASP protects alters chromatin accessibility. (A) Nucleosome profiles from HeLa cells stably transduced with empty vector (EV) or ASP-3xFLAG treated for 20 min with MNase at the indicated concentrations. MWM: molecular weight marker, U: unit. Mono, Di and Tri-nucleosome are highlighted. (B, C) Nucleosome intensity profiles of samples in (A) for the indicated MNase concentrations, normalized to the mono-nucleosome signal. Distance measures start at the bottom the gel (mono-nucleosome). (D) Ratio of poly-nucleosomes over mono- to tri-nucleosomes for cell lines form (A) at the indicated MNase concentrations.

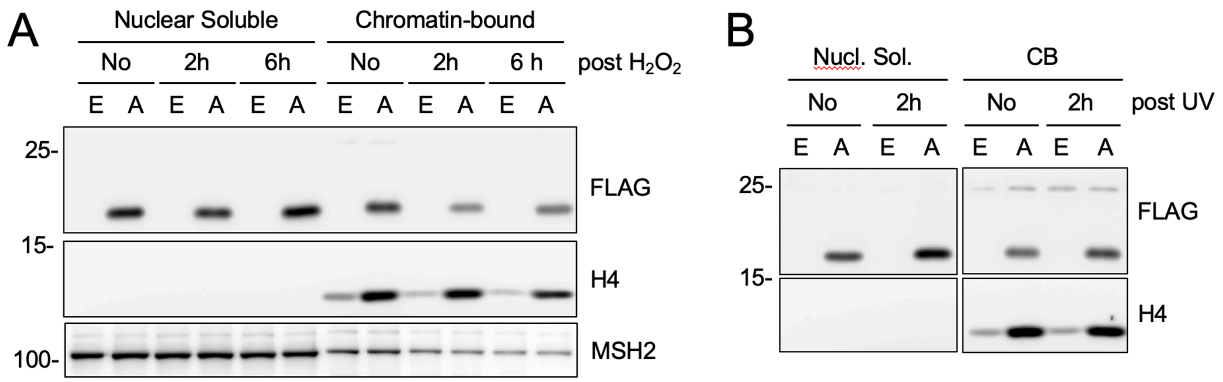


**Fig. S8.**

UV or $H_2O_2$ does not alter ASP chromatin association. (A) Expression of ASP-3xFLAG and the indicated proteins in soluble nuclear extract (NE) and chromatin bound protein (CB) from HeLa cells stably transduced with empty vector (EV) or ASP-3xFLAG. Cells were left untreated or collected at the indicated time points after $H_2O_2$ treatment (500 μM, 30 min), followed by Western blotting. (B) Western blot as in (A) in cells collected 2 h after treatment with 1 mJ UV. One of two independent experiments is shown.

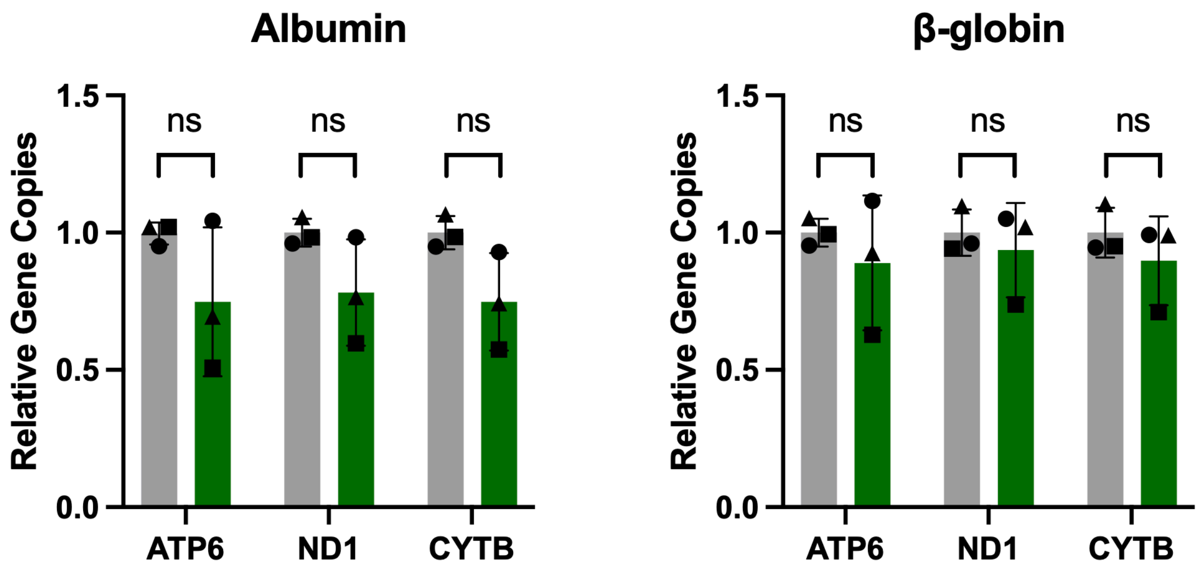


**Fig. S9.**

Quantitative PCR assays measuring cellular mtDNA copy number in HeLa-ASP vs. HeLa-EV cells. Copy number of three mtDNA-encoded genes (ATP6, ND1, Cytochrome B) normalized to copy numbers of 2 nDNA-encoded genes (albumin and b-globin; 2 copies/cell). No statistically significant difference was observed between the two cell lines.

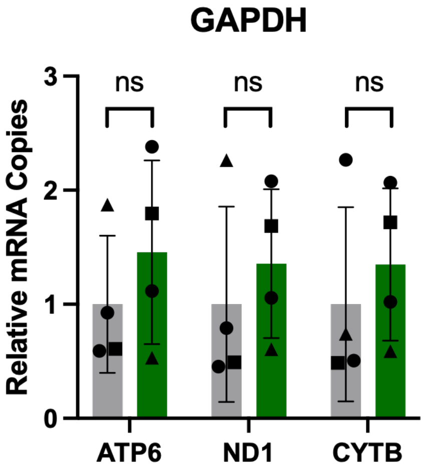


**Fig. S10.**

Quantitative RT-PCR assays measuring mtDNA transcription in HeLa-ASP vs. HeLa-EV cells. Copy number of mRNA of three mtDNA-encoded genes (ATP6, ND1, Cytochrome B) normalized to nDNA-encoded GAPDH mRNA copies. No statistically significant difference was observed between the two cell lines.

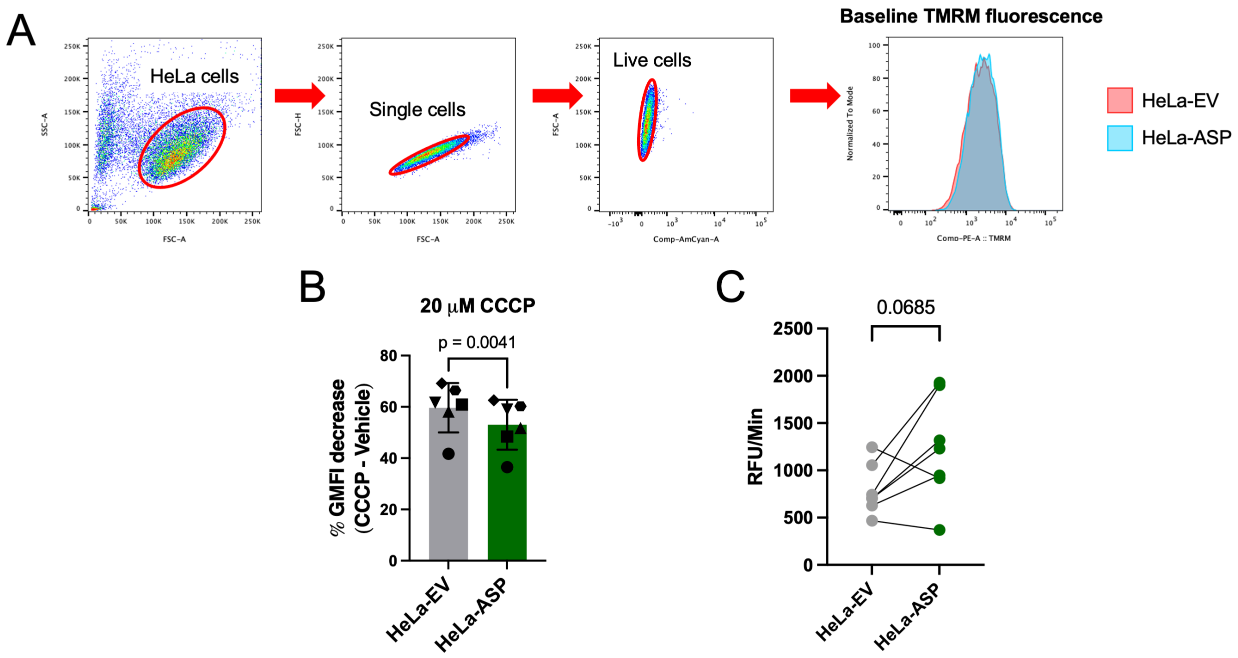


**Fig. S11.**

TMRM fluorescence analysis. (A) Gating strategy used for measurement of TMRM fluorescence as a function of mitochondrial membrane potential in HeLa-ASP and HeLa-EV cells. (B) Percent reduction of GMFI of TMRM as a function of the change in mitochondrial membrane potential following CCCP-induced depolarization (6 biological replicates). (C) Spare respiratory capacity ($OCR_{max} – OCR_{baseline}$; n = 7) of HeLa-ASP vs. HeLa-EV cells. Data show a trend toward greater spare respiratory capacity in ASP-expressing vs. control cells.

| **Antibody** | **Source** | **Catalog number** |
|---|---|---|
| 8-oxoG | Santa Cruz | sc-130914 |
| Anti-mouse-HRP | Cell Signaling Technology | 7076S |
| Anti-rabbit-HRP | Cell Signaling Technology | 7074S |
| ASP (324.6) | Raised in-house | See ref. 7 |
| ATP5A1 | Cell Signaling Technology | 18023S |
| ATP6 | Cell Signaling Technology | 70262S |
| ATP8 | Cell Signaling Technology | 17142S |
| b-Actin | Sigma Millipore | A3854 |
| CHD1 | Cell Signaling Technology | 4351S |
| COX1 | Cell Signaling Technology | 55159S |
| COX2 | Cell Signaling Technology | 31219S |
| COX4 | Cell Signaling Technology | 4850T |
| Cytochrome B | Cell Signaling Technology | 54618S |
| Cytochrome C | Cell Signaling Technology | 11940T |
| EPEA Tag (C-tag) | Thermo Fisher | MA562805 |
| FLAG (8H8L17) | Thermo Fisher | 701629M |
| FLAG (M2) | Millipore Sigma | F3165 |
| GAPDH | Cell Signaling Technology | 5174S |
| H2A | Cell Signaling Technology | 12349S |
| H2B | Cell Signaling Technology | 12364S |
| H3 | Cell Signaling Technology | 9715S |
| H4 | Cell Signaling Technology | 2935S |
| HIV-1 gp120 HRP | Thermo Fisher | PA1-73097 |
| HIV-1 p24 HRP | Thermo Fisher | PA1-73094 |
| HIV-1 Vpr | HIV Reagent Program | ARP 11836 |
| IPO9 | Bethyl | A305-475A |
| KPNA2 | Proteintech | 10819-1-AP |
| KPNB1 | Proteintech | 10077-1-AP |
| Lamin B1 | Proteintech | HRP-66095 |
| mH2A1.1 | Cell Signaling Technology | 12455S |
| mH2A1.2 | Sigma-Aldrich | MABE61 |
| Mouse IgG | Abcam | ab37355 |
| MPG | Proteintech | 11481-2AP |
| MRPL38 | Invitrogen | PAS-118074 |
| MRPL49 | Proteintech | 15542-1-AP |

| | | |
|---|---|---|
| MRPS35 | Proteintech | 16457-1-AP |
| MSH2 | Bethyl | A300-452A |
| MSH3 | Bethyl | A305-314A |
| ND3 | Cell Signaling Technology | 45859S |
| NDUFS1 | Cell Signaling Technology | 70264S |
| PARP1 | Proteintech | 13371-1-AP |
| PTCD3 | Proteintech | 25158-1-AP |
| Rabbit IgG | Abcam | ab37415 |
| RBBP5 | Cell Signaling Technology | 13171S |
| RISP | Cell Signaling Technology | 95231S |
| RPA32 | Thermo Fisher | MA1-26418 |
| RPA70 | Thermo Fisher | MA5-17166 |
| SSRP1 | Proteintech | 67313-A-Ig |
| SUPT16H | Cell Signaling Technology | 12191S |
| TNPO1 | Cell Signaling Technology | 31452S |
| Tubulin | Proteintech | 11224-1-AP |
| WDR5 | Cell Signaling Technology | 13105S |
| WDR82 | Thermo Fisher | PA5-83499 |
| XPC | Abcam | ab245385 |

**Table S1.**
List of antibodies used in this study.

| Primer | Sequence 5' - 3' |
| --- | --- |
| HIV-1 Sense F | AACTAGGGAACCCACTGCTTAAG |
| HIV-1 Sense R | TCTCCTTCTAGCCTCCGCTAGTC |
| HIV-1 Antisense RT | CTGATCTAGAGGTACCGGATCCAACATGTGGCAGGAAGTAGG |
| HIV-1 Antisense F | TGATGAACATCTAATTTGTCCACTGA |
| HIV-1 Antisense R | CTGATCTAGAGGTACCGGAT |
| ATP6 F | CACTAAAGGACGAACCTGATCTC |
| ATP6 R | GATAGTTGGGTGGTTGGTGTA |
| ND1 F | GAAGTCACCCTAGCCATCATTC |
| ND1 R | GCAGGAGTAATCAGAGGTGTTC |
| Cytochrome B F | ACCCTCACACGATTCTTTACC |
| Cytochrome B R | AGGGTGGAAGGTGATTTTATC |
| Albumin F | TGAAACATACGTTCCCAAAGAGTTT |
| Albumin R | CTCTCCTTCTCAGAAAGTGTGCATAT |
| β-globin F | CAACCTCAAACAGACACCATGG |
| β-globin R | TCCACGTTCACCTTGCCC |
| GAPDH F | CCAGCAAGAGCACAAGAGGA |
| GAPDH R | GAGATTCAGTGTGGTGGGGG |

**Table S2.**
List of primers used in this study.

**Data S1. (separate file)**
Complete list of 431 proteins significantly enriched (376 proteins) or depleted (55 proteins) in ASP-EPEA vs. ASP-WT samples.
